# HARP: Hierarchical Anatomical Refinement of Pathways for Whole-Brain Tractography

**DOI:** 10.64898/2026.05.27.728127

**Authors:** Simona Leserri, Kathleen S. Rockland, Dogu Baran Aydogan

## Abstract

Diffusion MRI tractography is an ill-posed inverse problem that requires anatomical constraints to ensure the plausibility of reconstructed white-matter pathways. Yet, encoding whole-brain constraints is challenging because neuroanatomical knowledge is fragmented and unevenly distributed across regions and spatial scales. We introduce HARP, a flexible framework that hierarchically injects anatomical constraints at different levels of detail, in parallel with more and more granular brain segmentations. Unlike existing fixed rule-based approaches, HARP enables the systematic, scalable integration of diverse priors within a unified framework. Across multiple tractography algorithms and acquisition protocols, HARP achieves up to a 9% reduction in implausible streamlines. These rejected connections are absent from a ground truth brain phantom, indicating their artifactual origin, and cannot be reliably identified using post-hoc filtering weights alone. By reducing false positive reconstructions, HARP improves the anatomical specificity of tract reconstructions and downstream connectome estimates. More broadly, HARP represents a step toward a collaborative effort to encode neuroanatomical insights in tractography pipelines, with the goal of advancing in vivo whole-brain connectivity studies.

## 1. Introduction

Understanding the architecture of the brain’s white matter is a major scientific endeavor (Schmahmann & Pandya, 2006), as it constitutes the physical substrate for connections and information transfer between areas. White-matter connectivity is altered in several diseases such as stroke, epilepsy, and neurodegenerative cognitive disorders (Ciccarelli et al., 2008). Currently, diffusion magnetic resonance imaging (dMRI)-based tractography is the only widely available technique for studying whole-brain structural connectivity in vivo and non-invasively. Tractography is useful in many clinical applications (Yamada et al., 2009), including neurosurgical planning (Essayed et al., 2017) and brain stimulation studies (Calabrese, 2016; Mayberg, 2009).

While indispensable for mapping the brain’s structural connectivity, tractography lacks the anatomical precision of invasive ex vivo techniques (Thomas et al., 2014). Tractography estimates long-range streamlines from local dMRI-inferred fiber orientations (Jeurissen et al., 2019). It is therefore inherently an inverse, ill-posed problem: inverse in that, from known dMRI data, it estimates unknown fiber trajectories; ill-posed in that multiple fiber distributions can equally explain the same diffusion signal (Daducci et al., 2025; Jeurissen et al., 2019). Several well-known tractography limitations—including false positive (Maier-Hein et al., 2017; Thomas et al., 2014) and false negative (Aydogan et al., 2018) reconstructions, challenges in resolving crossing fibers (Jeurissen et al., 2019; K. Schilling et al., 2017), the gyral bias (K. Schilling et al., 2018), and the wall, bottleneck and narrow intersection effects (Rheault et al., 2025) — stem from tractography being an inverse, ill-posed problem. Moreover, the choice of a tractography algorithm and its parameters strongly affect the resulting tractogram (Aydogan et al., 2018). Local tractography methods, which perform stepwise integration of the local orientation information to independently propagate each streamline, are the most widely used (Jeurissen et al., 2019). In contrast, global tractography methods seek a set of trajectories in the whole brain that jointly explain the diffusion signal (Daducci et al., 2025; Jeurissen et al., 2019).

In this study, we focus on local tractography methods and on the pervasive issue of false positive streamlines, which are tractography-based connections obtained from dMRI data without an anatomical counterpart. In an international tractography challenge, these invalid connections were, on average, reported to be four times more likely to be reconstructed than valid ones (Maier-Hein et al., 2017). The problem is more pronounced in lower-quality acquisitions, such as those routinely obtained in clinical settings (Dell’Acqua et al., 2025; Maier-Hein et al., 2017; Thomas et al., 2014). However, even high-quality scans, for example those with increased image resolution (K. G. Schilling et al., 2019), are unlikely to fully eliminate false positives since crossings and ambiguous fiber configurations persist and lead to spurious streamlines (K. Schilling et al., 2017).

Given that the ill-posed nature of the problem is a key factor behind tractography errors, specificity can be improved using explicit priors and constraints to steer reconstructions toward anatomically plausible tractograms. Several such strategies have been developed, including the application of geometric constraints, anatomical priors, or post-hoc filtering. Geometric constraints such as minimum or maximum length, curvature/angle thresholds, and step-size controls are routinely enforced during tracking to reject physically implausible paths (Jeurissen et al., 2019).

Anatomically Constrained Tractography (ACT) applies explicit anatomical priors (R. E. Smith et al., 2012). ACT integrates tissue-specific information to guide whole-brain streamline generation and termination according to the following rules: streamlines crossing the cerebrospinal fluid are rejected; in white matter, terminations are prohibited unless exiting the tracking mask (e.g., toward the spinal cord); terminations are valid in cortical and subcortical gray matter, but once a streamline enters subcortical gray matter, it cannot re-enter other tissues. Similarly, ExTractorFlow (Petit et al., 2019, 2023) filters out streamlines crossing the cerebrospinal fluid and those ending in the white matter. Its sequential pipeline also removes streamlines below a minimum length, those completing a 360° turn, and those crossing the sulci. Such anatomical-rule-based strategies can substantially reduce false positive connections and improve anatomical plausibility, including in lower-quality data, but they do not fully resolve false positive reconstructions (Maier-Hein et al., 2017; R. E. Smith et al., 2012; Yeh et al., 2016) .

A complementary strategy to reduce false positive streamlines is to apply post-hoc filters. Filtering techniques attach quantitative information to tractograms by assigning streamline weights that match either the dMRI signal, as in Convex Optimization Modeling for Microstructure Informed Tractography (COMMIT2) (Schiavi et al., 2020), or the local fiber orientation distribution (FOD), as in spherical-deconvolution informed filtering of tractograms (SIFT2) (R. E. Smith et al., 2015). In COMMIT2, weights are estimated by fitting a microstructural local forward model to the diffusion signal. Additionally, based on their connectivity, streamlines are grouped into bundles, enforcing anatomical organization (Bullmore & Sporns, 2012). Despite their different mechanisms, both algorithms assign low weights to unlikely connections. However, because these weights are primarily derived based on the diffusion signal or FOD, they do not explicitly consider the anatomical plausibility of streamlines. Moreover, their performance worsen as fiber complexity increases, motivating further improvements grounded in neuroanatomical knowledge (Sarwar et al., 2023).

A wealth of detailed anatomical knowledge is available, but much of it remains difficult to incorporate systematically into tractography. High-resolution insights from dissection and tracing studies are challenging to translate into human whole-brain tractography rules because they are fragmented across regions, spatial scales, and species (Descoteaux et al., 2025). Notably, the categorical classification of cortical connections into associations, projections, and commissural fibers (Catani, 2025; Meynert & Sachs, 1885; Schmahmann & Pandya, 2006) is not yet routinely incorporated when obtaining whole-brain tractograms, with some exceptions such as (Petit et al., 2023). Moreover, consistent with work allowing streamlines to traverse subcortical regions (Behrens, Johansen-Berg, et al., 2003; Draganski et al., 2008; Palesi et al., 2015), modifications to the subcortical ACT rule have been suggested (Bajada et al., 2019). Complex areas such as the brainstem may also require additional rules that are not currently leveraged to improve tractograms.

Overall, a method that hierarchically injects increasingly precise, region-informed constraints would be highly useful to improve whole-brain tractogram specificity, without discarding plausible connections. To that end, we propose a new strategy called Hierarchical Anatomical Refinement of Pathways (HARP). HARP builds upon and extends the ACT framework in a hierarchical way, enabling progressive refinement of connectivity estimates as additional anatomical priors become available, particularly through increasingly precise segmentations of brain regions.

Preliminary HARP results were presented at the 2024 annual meeting of the International Society for Magnetic Resonance in Medicine (Leserri & Aydogan, 2024). In this study, we provide a complete description of HARP and comprehensively evaluate its ability to increase tractogram specificity and anatomical plausibility. Through HARP, we identify artefactual, anatomically implausible trajectories reconstructed by tractography from an anatomically curated ground truth brain phantom. We quantify reduction in implausible streamlines across tractography algorithms and clinical acquisition protocols as well as in high-quality research data. Finally, we compare streamline-level weights assigned by SIFT2 and COMMIT2 and evaluate how consistent these techniques score the streamlines that are identified as implausible by HARP.

## 2. Methods

### 2.1 Hierarchical anatomical segmentation

HARP uses anatomical regions of interest (ROIs) at progressively finer levels of detail (see Fig. 1). Then, it applies anatomical constraints to tractograms following the hierarchical segmentation levels. Each level refines the set of “plausible” streamlines identified at the previous level, reclassifying some as “implausible” based on the more detailed segmentation and rules. Plausible streamlines are classified based on their endpoints or the regions they traverse.

**Figure 1.**
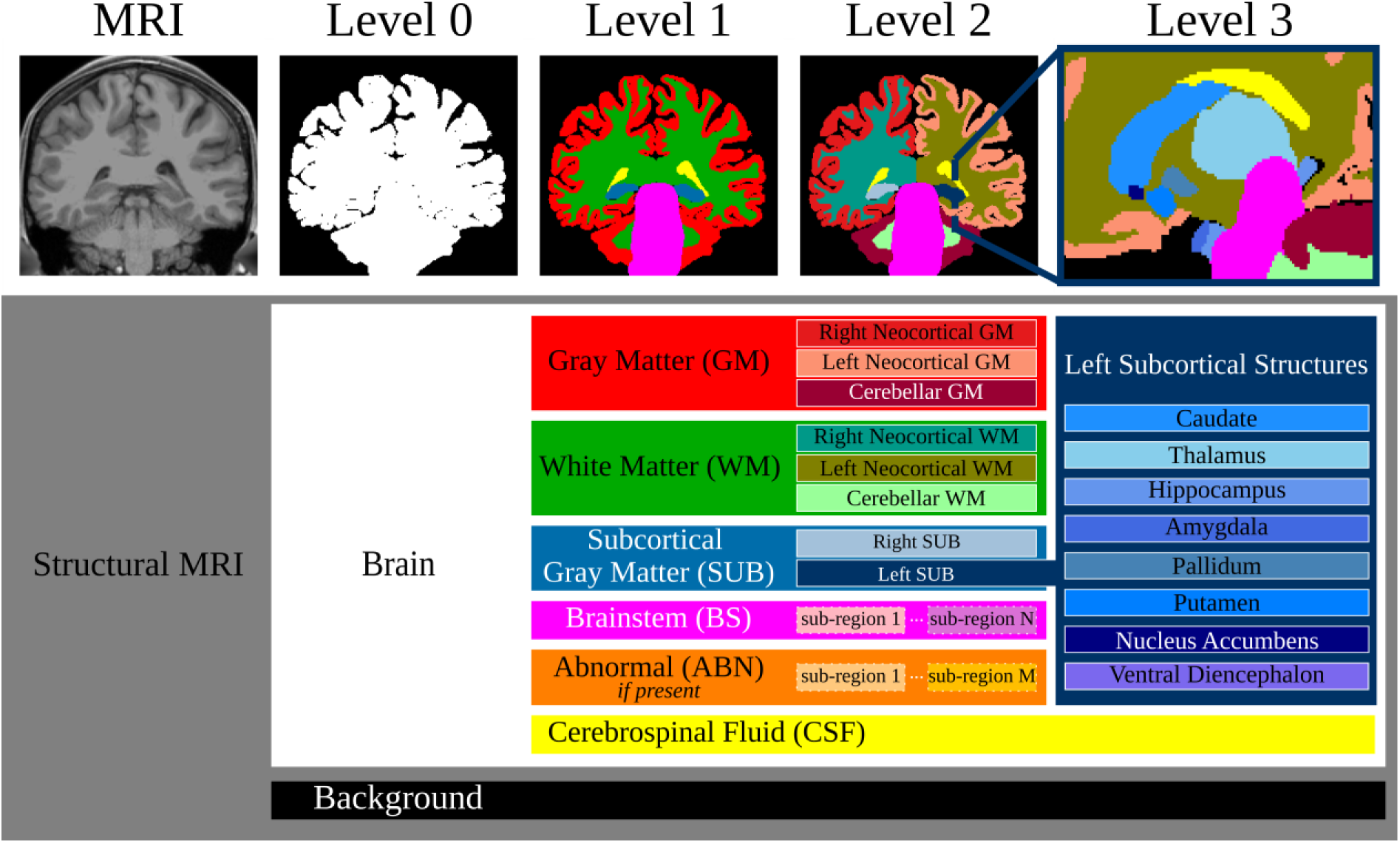
Hierarchical anatomical segmentation. Illustration of a multi-level segmentation framework applied to structural brain MRI data. The top panel shows the hierarchical segmentation of brain regions. The bottom panel shows how regions are organized across hierarchical levels. At level 0, the brain is separated from the non-brain area. At level 1, the brain is segmented into six main classes: Gray Matter (GM), White Matter (WM), Subcortical Gray Matter (SUB), Brainstem (BS), Cerebrospinal Fluid (CSF), and Abnormalities (ABN), which includes pathological tissues when present. At level 2, level 1 regions are refined into smaller subregions. GM comprises Left Neocortical GM (LN_GM), Right Neocortical GM (RN_GM), and Cerebellar GM (CER_GM). WM comprises Left Neocortical WM (LN_WM), Right Neocortical WM (RN_WM), and Cerebellar WM (CER_WM). SUB comprises Left Subcortex (L_SUB) and Right Subcortex (R_SUB). BS and ABN remain unsegmented in our implementation, yet further segmentation is possible. At level 3, finer grained segmentations are introduced. In the case of L_SUB structures, specific nuclei such as the Caudate, Thalamus, Hippocampus, Amygdala, Pallidum, Putamen, Nucleus Accumbens, and Ventral Diencephalon are identified. Color coding reflects class and segmentation level.

### 2.2 Rules to refine pathways

Below are the rules and classification logic used in HARP (see Fig. 2).

**Figure 2.**
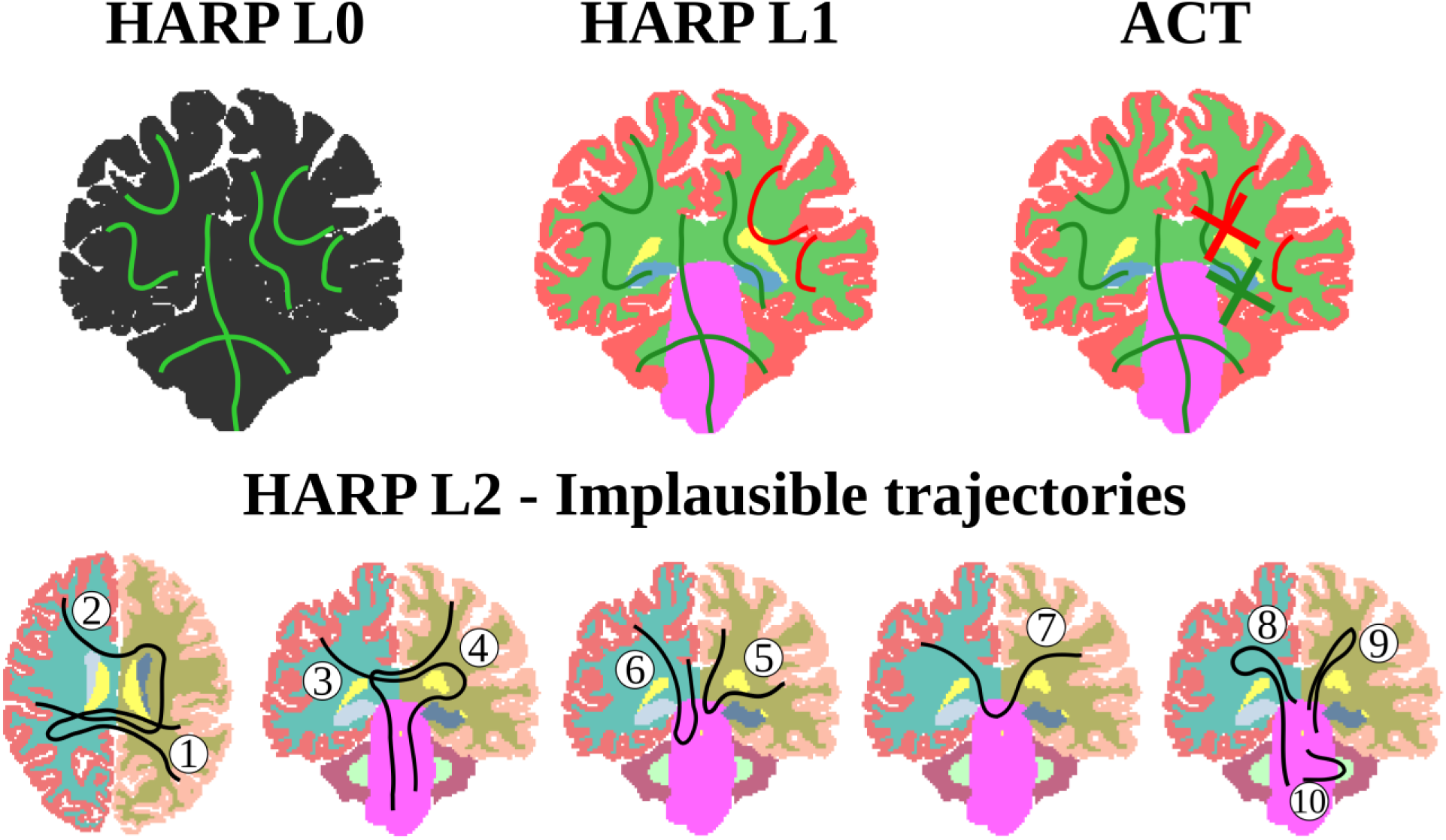
Anatomical rules of HARP levels. Illustrative example drawings show plausible trajectories in green and implausible ones in red or black, according to the constraints defined for each HARP level. The background structural image is displayed using the same color scheme as the corresponding segmentation levels in Figure 1. Streamline endpoints determined during fiber tracking with ACT are indicated by crosses. Numbers associated with level 2 (L2) streamlines indicate the specific implausible trajectory types defined in Section 2.2. Rules to refine pathways – level 2.

#### LEVEL 0 (L0)

- Based on 1 ROI: Brain mask
- RULE 1: Streamlines must be entirely defined inside the brain mask, or L0 ROI. Upon leaving such mask, the streamline is terminated and accepted, allowing for instance streamlines to travel towards the spinal cord.
- Classification: Non applicable as all generated streamlines are considered plausible at this level.

#### LEVEL 1 (L1)

This level is equivalent to traditional ACT (R. E. Smith et al., 2012), with rules being dependent on tissue types. We define as endpoint ROI those in which streamlines are allowed to terminate.

- Based on 6 ROIs (4 of which are endpoints *): Gray Matter (GM)*, White Matter (WM), Subcortical Gray Matter (SUB)*, Brainstem (BS)*, Cerebrospinal Fluid (CSF), and Abnormalities (ABN)*
- Rules:
  - RULE 2: No crossing of CSF
  - RULE 3: Streamline endpoint in any endpoint ROIs
- Classification: 10 end-to-end connectivity combinations (6 if ABN is not available) for plausible streamlines; 3 combinations (Not satisfying both rule 2 and rule 3, not satisfying only rule 2, not satisfying only rule 3) for implausible streamlines.

#### LEVEL 2 (L2)

- Based on 11 ROIs (7 of which are endpoints*): Left Neocortical GM (LN_GM)*, Right Neocortical GM (RN_GM)*, Cerebellar GM (CER_GM)*, Left Neocortical WM (LN_WM), Right Neocortical WM (RN_WM), Cerebellar WM (CER_WM), Left Subcortex (L_SUB)* and Right Subcortex (R_SUB)*, BS*, CSF, ABN*.
- Rules: Currently, these rules enforce categorical separation of cortical connections into three groups following (Catani, 2025): associations, where both streamline endpoints lie within the same cortical hemisphere; commissural fibers, where endpoints lie in opposite hemispheres; and projections, where one endpoint is in either cortical hemisphere and the other in the brainstem. Streamlines starting and ending in the brainstem and taking long courses through other areas are also discarded.
  - RULE 4 (trj1): No right associations crossing the left hemisphere
  - RULE 5 (trj2): No left associations crossing the right hemisphere
  - RULE 6 (trj3): No left projections crossing the right hemisphere
  - RULE 7 (trj4): No right projections crossing the left hemisphere
  - RULE 8 (trj5): No right associations crossing the brainstem
  - RULE 9 (trj6): No left associations crossing the brainstem
  - RULE 10 (trj7): No commissural fibers crossing the brainstem
  - RULE 11 (trj8): No brainstem-to-brainstem connections crossing left cortical white matter
  - RULE 12 (trj9): No brainstem-to-brainstem connections crossing right cortical white matter
  - RULE 13 (trj10): No brainstem-to-brainstem connections crossing cerebellar white matter
- Classification: 28 end-to-end connectivity combinations (21 if ABN is not available) for plausible streamlines; 10 trajectories for implausible streamlines.

#### FURTHER LEVELS

With more detailed segmentations, additional HARP levels can introduce new anatomical rules tailored to finer subdivisions, such as specific subcortical nuclei. At these levels, implausible trajectories could be identified based on known neuroanatomical pathways. These increasingly specialized constraints would further narrow the set of plausible streamlines while maintaining the hierarchical refinement principle of HARP.

### 2.3 Implementation

In our current implementation, for the ROIs, we adopted surfaces instead of images to remove spatial mismatches due to partial volume effects, similarly to (Warrington et al., 2020; Yeh et al., 2017). To that end, in our open-source software Trekker (https://dmritrekker.github.io/), we implemented *prepXact*, a command that generates high-resolution surface meshes starting from segmentation of a structural T1-weighted image through a marching cubes algorithm.

Taking as input these combined surfaces and any tractogram, the *harp* command, part of Trekker (available in harp branch), evaluates each streamline against 27 criteria (see Appendix -Criteria), yielding a sparse binary #streamlines × #criteria table.

Through logical operations on these columns, our code then checks for the 13 HARP rules and flags each streamline as being L1 plausible or implausible, L2 plausible or implausible, and determines L1 and L2 classification. These rules being independent, classification may not be unique, for instance because of overlapping ROIs. Ambiguous streamlines are assigned to a unique class by means of priority list defined in our code (see 3.4 and Appendix – Priority order). For each streamline, the command returns labels for L1 and L2 classification; L1 and L2 implausible classification; and L1 and L2 (im)plausibility and ambiguity flags. These labels and flags are used to select the subset of streamlines of interest from the input tractogram.

## 3. Experiments

### 3.1 Data

#### 3.1.1 ISMRM 2015 Challenge Dataset

We used the ISMRM 2015 Challenge dataset (http://www.tractometer.org/ismrm_2015_challenge/data) (Maier-Hein et al., 2017) for qualitative demonstration only. It consists of a synthetic whole-brain dMRI dataset generated from a manually curated selection of 25 major fiber bundles of HCP subject 100307. Thus, this dataset represents a controlled setting for visually assessing the generation of HARP level 2 implausible trajectories, as the curated bundles themselves do not contain any such trajectories.

The dMRI data features a voxel dimension of 2×2×2 mm^3^, 32 gradient directions, and a b-value of 1000 s/mm2, supplemented by two b0 images with reverse phase encodings. To prepare the data, we applied the MPPCA algorithm (Veraart, Fieremans, et al., 2016; Veraart, Novikov, et al., 2016) for denoising and removed Gibbs ringing artifacts using the method described in (Kellner et al., 2016). Geometric distortions were corrected using FSL’s *topup* (Andersson et al., 2003) and eddy_*openmp* (Andersson & Sotiropoulos, 2016; S. M. Smith et al., 2004). Finally, Fiber Orientation Distributions (FODs) were computed using the algorithm in (Tran & Shi, 2015) and represented via coefficients of 8^th^ order spherical harmonics.

#### 3.1.2 TractoInferno Cohorts

We used the publicly available TractoInferno clinical dataset (Poulin et al., 2022), which includes data of 354 healthy participants unevenly distributed across six cohorts. For 284 of those subjects, a T1-weighted image, a preprocessed diffusion-weighted image, and FODs of order 8 are accessible in OpenNeuro (https://openneuro.org/datasets/ds003900/versions/1.1.1).

#### 3.1.3 HCP Cohort

We additionally included preprocessed dMRI data from the HCP Q1 release (Glasser et al., 2013; Van Essen et al., 2012), acquired at 1.25-mm isotropic resolution with three shells (b = 1000, 2000, 3000 s/mm²), 90 directions per shell, and two-phase encoding directions. FODs of order 8 were estimated following (Tran & Shi, 2015).

#### 3.1.4 Subject Selection Criteria

For each cohort, we retained 14 subjects. This corresponds to including all subjects from the smallest TractoInferno cohort and, for every remaining cohort, selecting the first 14 subjects in numerical order of subject ID for which all surface processing steps, tractography, and quality control checks were successful. We thus included 98 subjects in the quantitative analyses.

### 3.2 Surface processing

All cohorts underwent identical surface processing. We applied FreeSurfer’s (Fischl, 2012) *recon-all* pipeline to generate anatomical segmentations and parcellations. Using Trekker’s *prepXact* command, we extracted high-resolution surface meshes for the following anatomical regions: WM (further segmented into L_WM, R_WM, CER_WM), GM (further segmented into LN_GM, RN_GM, CER_GM), SUB (further segmented into L_SUB, R_SUB), CSF, BS. Their union was converted to an image and used as the L0 ROI. No ABN surface was identified in our healthy population.

### 3.3 Tractography

For each TractoInferno and HCP subject, four tractography algorithms were used:

a. deterministic SD_STREAM (J. D. Tournier et al., 2012) with default parameters,
b. probabilistic iFOD2 (J. Tournier et al., 2010) with default parameters,
c. Parallel Transport Tractography (PTT) (Aydogan & Shi, 2021) with default parameters, and minDataSupport=0.025, and dataSupportExponent=0.5, and
d. Modified PTT, which is same as (c) but with modified parameter values of minDataSupport=0.005, minRadiusOfCurvature=0.5 mm.

For SD_STREAM and iFOD2, MrTrix3 was used (J. D. Tournier et al., 2019). For PTT, Trekker was used (Aydogan & Shi, 2021).

For the qualitative demonstration in the ISMRM 2015 challenge dataset, we only used the modified PTT approach.

For all algorithms, seeding was performed inside the L0_ROI mask, and 10M streamlines of lengths between 2.5 mm and 250 mm were generated. The length histogram of each of the resulting 4*98 tractograms was visually inspected for quality control.

### 3.4 HARP

With the tractograms and the ROIs surfaces as input, we run HARP. We considered L1 plausible the streamlines satisfying HARP rules 1, 2, and 3. The L1 plausible tractograms contain exclusively L1 plausible streamlines. We considered L2 plausible the streamlines satisfying the above rules and NOT matching the rules 4 to 13 for the implausible trajectories. The L2 plausible tractograms contain exclusively L2 plausible streamlines. Additionally, the endpoint surfaces allow for L1 and L2 endpoint categorization of each streamline. In case of endpoint ROI surface intersection, the classification of the streamline was marked as ambiguous. To resolve the ambiguity, and ensure that each streamline had a unique classification, we applied the priority order provided in the Appendix.

### 3.5 HARP-SIFT2-COMMIT2 comparison

The filtering algorithms were compared on L1 plausible tractograms generated with the default PTT tracking configuration of the TractoInferno and HCP cohort. We chose L1 plausible tractograms because they closely resemble ACT, and the default PTT method as it yielded the fewest number of L2 implausible streamlines (see Fig. 5). Because COMMIT2 incorporates anatomical priors by grouping streamlines whose endpoints lie within the same parcel, we further restricted the analysis to streamlines whose endpoints fell in parcels of the Desikan–Killiany atlas (Desikan et al., 2006) obtained with FreeSurfer. These connecting L1 plausible streamlines, for which HARP categorization was already available, were used as input to both filtering algorithms.

We ran COMMIT2 (v2.4.2) using a Stick-and-Ball forward model (Behrens, Woolrich, et al., 2003): the stick compartment modeled intra-axonal diffusion with parallel diffusivity 1.7 × 10−3 mm²/s and zero perpendicular diffusivity; the isotropic (ball) compartments modeled extra-axonal tissue with diffusivity 1.7 × 10−3 mm²/s and free water with diffusivity 3.0 × 10−3 mm²/s. Because there is no closed-form solution for the COMMIT2 group-sparsity regularization parameter (λ), we empirically tested λ ∈ [10−7, 10−1] and selected λ = 4×10−4 as the largest value for which the median matrix density reduction across subjects remained <50%. We ran SIFT2 (MRtrix3 v3.0.8) with default parameters.

For each subject and for each streamline, we recorded the HARP L2 implausible flag, the SIFT2 weight, and the COMMIT2 weight. To obtain a consistent subsample of streamlines across subjects, we first generated a reproducible set of indices by shuffling the integers from 1 to 1,000,000 using a fixed random seed and selecting the first 100,000 unique values. For each subject, the streamlines corresponding to these indices were extracted from the L1 plausible tractograms. We compared HARP, SIFT2 and COMMIT2 using this matched subset of 100,000*98 streamlines.

Because SIFT2 and COMMIT2 assign streamline weights using fundamentally different modeling principles, their resulting weight distributions are not directly comparable. COMMIT2 explicitly promotes sparsity in the connectome, whereas SIFT2 produces a continuous, right-skewed distribution of weights. To allow fair cross-method comparisons, we harmonized the weights by ranking streamlines within each method and using the rank as a normalized weight measure. This approach preserves within-method ordering while eliminating scale differences that would otherwise bias comparisons.

### 3.6 Evaluation and statistics

To quantitatively estimate the impact of HARP on tractograms across cohorts and tracking algorithms, we performed several analyses. Since HARP level 1 rules closely mirror the widely adopted and established ACT rules, we mainly focus on the effect of additionally incorporating HARP level 2 rules.

1. We quantified the ratio of L1 implausible over L0 generated streamlines (all plausible); and the ratio of L2 implausible over L1 plausible streamlines. With the ratio implausible/plausible being our metric of interest, we ask: Are there differences among the 7 cohorts in the ratio of implausible/plausible streamlines? Are there differences among the 4 algorithms in the ratio of implausible/plausible streamlines? Do algorithm differences depend on cohort (Cohort × Algorithm interaction)? To answer these questions in a single framework, for each ratio we fit a separate linear mixed-effects model with Cohort (7 levels), Algorithm (4 levels), and their interaction (Cohort × Algorithm) as fixed effects, and a random intercept for Subject to account for repeated measurements. We used sum-to-zero (effect) coding for Cohort and Algorithm to obtain reference-free estimates and ANOVA-style tests of factors. Because the dependent variable is a proportion bounded between 0 and 100, values were rescaled to the (0,1) interval and analyzed on the logit scale, which provides variance stabilization. The model was estimated by restricted maximum likelihood (REML) as implemented in statsmodels (v0.14.6) (Seabold & Perktold, 2010). Model adequacy was evaluated through visual inspection of residual–fitted plots, histograms of residuals, and quantile–quantile plots. Convergence diagnostics and random effects variance estimates were examined to verify model stability. Inference on fixed effects was based on Wald tests with model-based covariance estimates. Post-hoc pairwise comparisons included simple-effects tests and model-based estimated marginal means, with Holm correction applied to control the family-wise error rate (Holm, 1979).
2. The L1 plausible streamlines contain both L2 plausible and L2 implausible streamlines. For each group, we quantified their endpoint distribution with respect to the three L1 endpoint ROIs, yielding two 3×3 triangular endpoint–endpoint matrices (twelve elements). Because no rules were imposed for subcortical structures, three of these elements were structurally zeros. The remaining nine elements form a valid composition and were renormalized to sum to 1. To assess whether the method, the cohort, or their interaction significantly affected the overall composition, we applied compositional data analysis using an isometric log-ratio (ilr) transformation, followed by a linear mixed-effects models for each composition coordinate with Subject as a random factor and Method, Cohort, and their interaction as fixed effects. Significance of fixed effects was evaluated using Wald tests, and p-values across ilr coordinates were combined according to the Fisher’s method and Holm corrected for multiple comparisons.
3. We examined whether the binary HARP L2 implausible categorization aligned preferentially with higher harmonized weights from SIFT2, COMMIT2, or both. For each filtering method, we computed A) the histogram of harmonized weights for the L2 implausible streamlines and B) the empirical cumulative distribution function (ECDF) of harmonized weights for the L2 implausible streamlines. We compared the weights using all streamlines pooled together and stratified by cohort.

## 4. Results

### 4.1 Qualitative

Figure 3 qualitatively illustrates the detection of implausible trajectories by HARP level 2 using the ISMRM 2015 challenge dataset. Comparison between the anatomically curated bundles used to generate the phantom (left) and the tractography-derived streamlines (right) reveals the presence of trajectories that are absent from the ground truth. These streamlines fall into HARP level 2 rules and are thus identified as false positive connections introduced by tractography.

**Figure 3.**
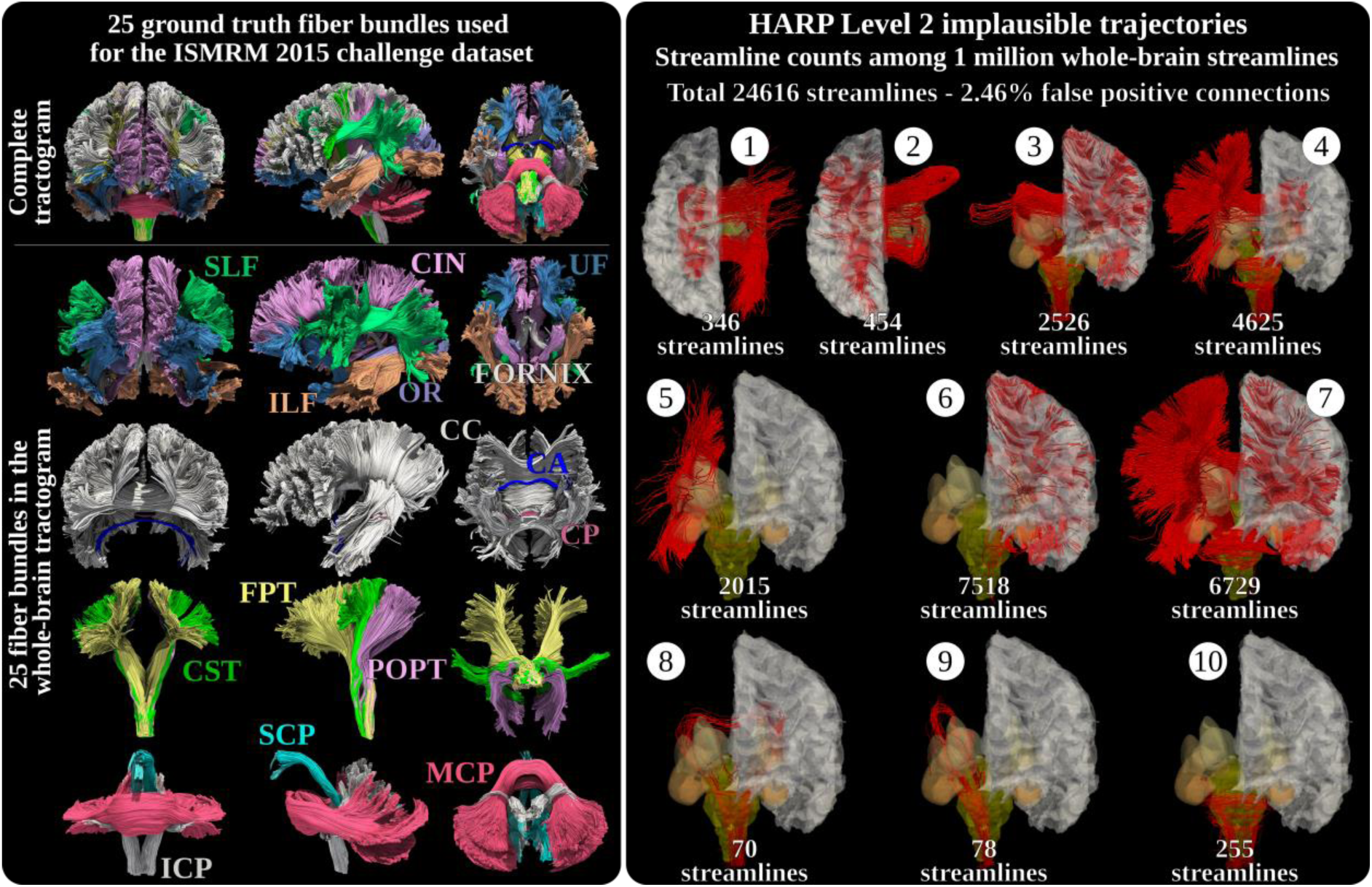
Qualitative demonstration of HARP level 2 implausible trajectories obtained using the ISMRM 2015 challenge dataset. The figure shows, on the left, the original whole-brain fiber bundles used for the generation of the phantom. SLF: Superior Longitudinal fasciculus; CIN: Cingulum; UF: Uncinate Fasciculus; Fornix; ILF: Inferior Longitudinal Fasciculus; OR: Optic Radiation; CC: Corpus Callosum; CA: Anterior Commissure; CP: Posterior Commissure; CST: cortico-spinal tract; FPT: frontopontine tract; POPT: parieto-occipital pontine tract; ICP: inferior cerebellar peduncle; SCP: superior cerebellar peduncle; MCP: middle cerebellar peduncle. These manually curated streamlines do not contain the implausible connections generated through tractography shown in the right panel. Overall, in this case, HARP identified 2.46% false positive streamlines based on level 2 rules. Numbers associated with level 2 (L2) streamlines in the right panel match those in Figure 2 and the implausible trajectory types defined in Section 2.2. Rules to refine pathways – level 2.

Figure 4 qualitatively shows the effect of HARP on the Default PTT tractogram of a TractoInferno subject. After application of level 1 rules, closely related to the commonly used ACT rules, tractograms contain implausible streamlines, in red in the top row of Figure 4. Adopting Harp level 2 rules yields cleaner and distinct associations, projections, and commissural fibers.

**Figure 4.**
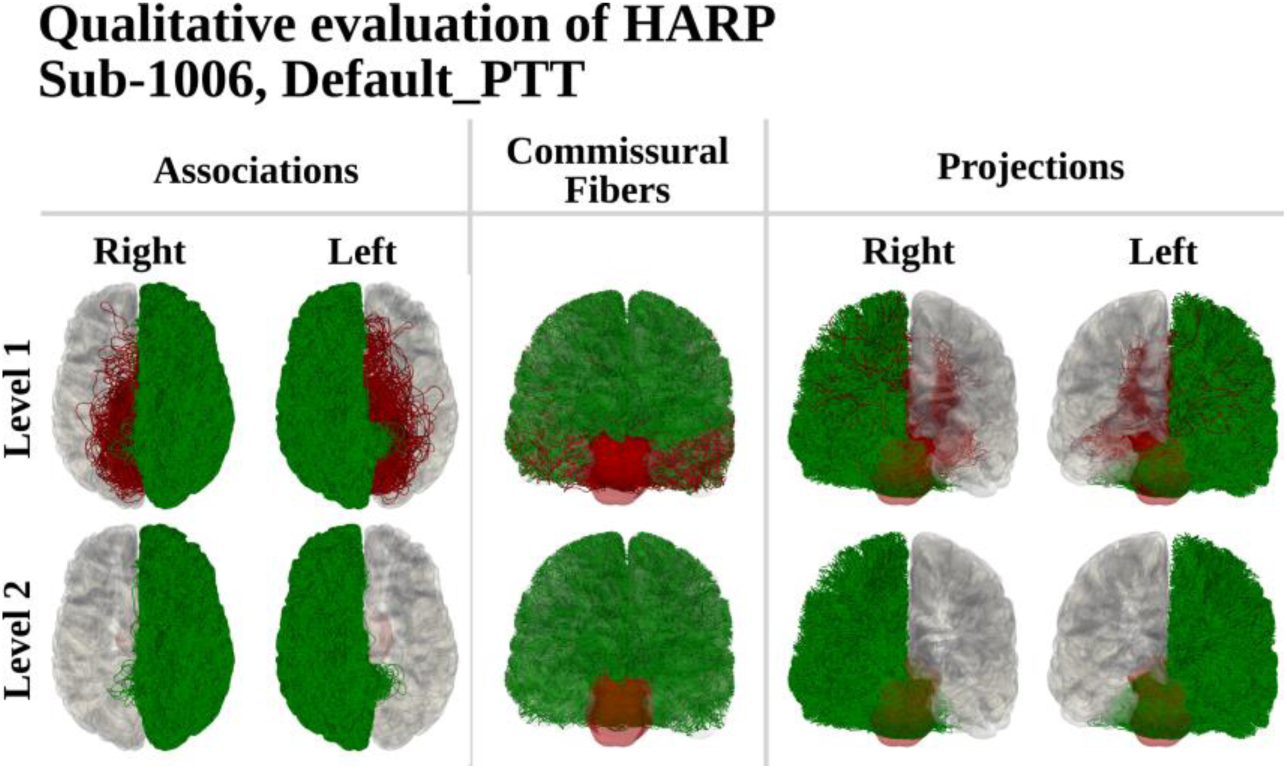
Qualitative effect of HARP on whole-brain tractography. The figure illustrates the impact of applying HARP rules to the tractogram of a representative TractoInferno subject obtained with the Default PTT algorithm. Using HARP level 1 results in implausible streamlines (shown in red in the top row), many of which are typically retained in tractograms and consequently corrupt downstream analyses. Enforcing HARP level 2 yields cleaner and well-differentiated associations, projections, and commissural pathways. Surfaces were obtained with FreeSurfer; gray matter is rendered in gray and the brainstem in light red, both semi-transparent. Association pathways are displayed in axial view, while projection and commissural fibers are shown in coronal view.

### 4.2 Quantitative

By applying the defined priority order, we uniquely classified all ambiguous streamlines. On average, across all cohorts and algorithms, these cases occurred among 2.71±2.38% of the L2 plausible streamlines and 4.18±2.46% among the L2 implausible streamlines.

Figure 5 quantifies the ratio of L1 implausible/L0 plausible streamlines across cohorts and tracking algorithms (top panel); and the ratio of L2 implausible/L1 plausible streamlines across cohorts and tracking algorithms (bottom panel). HARP level 1 rules flag as implausible between 55.90% and 93.50% of L0 streamlines. The logit-transformed ratio of L1 implausible to L0 plausible streamlines was modeled using a linear mixed-effects model (log-likelihood = 236.9032; random-intercept variance = 0.004 ± 0.012). Diagnostic plots indicated good model fit (Supplementary Figure 1). Wald tests identified significant main effects of Cohort, Algorithm, and their interaction. Estimated marginal means further indicated significant differences among all algorithm pairs and among 16/21 cohort pairs. Post-hoc simple-effects analyses showed significant Algorithm differences in 33/42 cohort-specific contrasts and significant Cohort differences in 35/84 algorithm-specific contrasts. In particular, within the iFOD2 algorithm, differences were significant in all pairwise cohort comparisons but for Mazoyer – Tamm, Poldrack – Tremblay, Tamm – Tsushida, Mazoyer – Tsushida; within the SD_STREAM algorithm, all pairwise post-hoc differences with the HCP cohort were statistically significant, and other comparisons were heterogeneous; within the Default PTT algorithm, only differences between the HCP and the clinical cohorts resulted significant; and within the Modified PTT algorithm, no significant difference between cohorts was identified.

**Figure 5.**
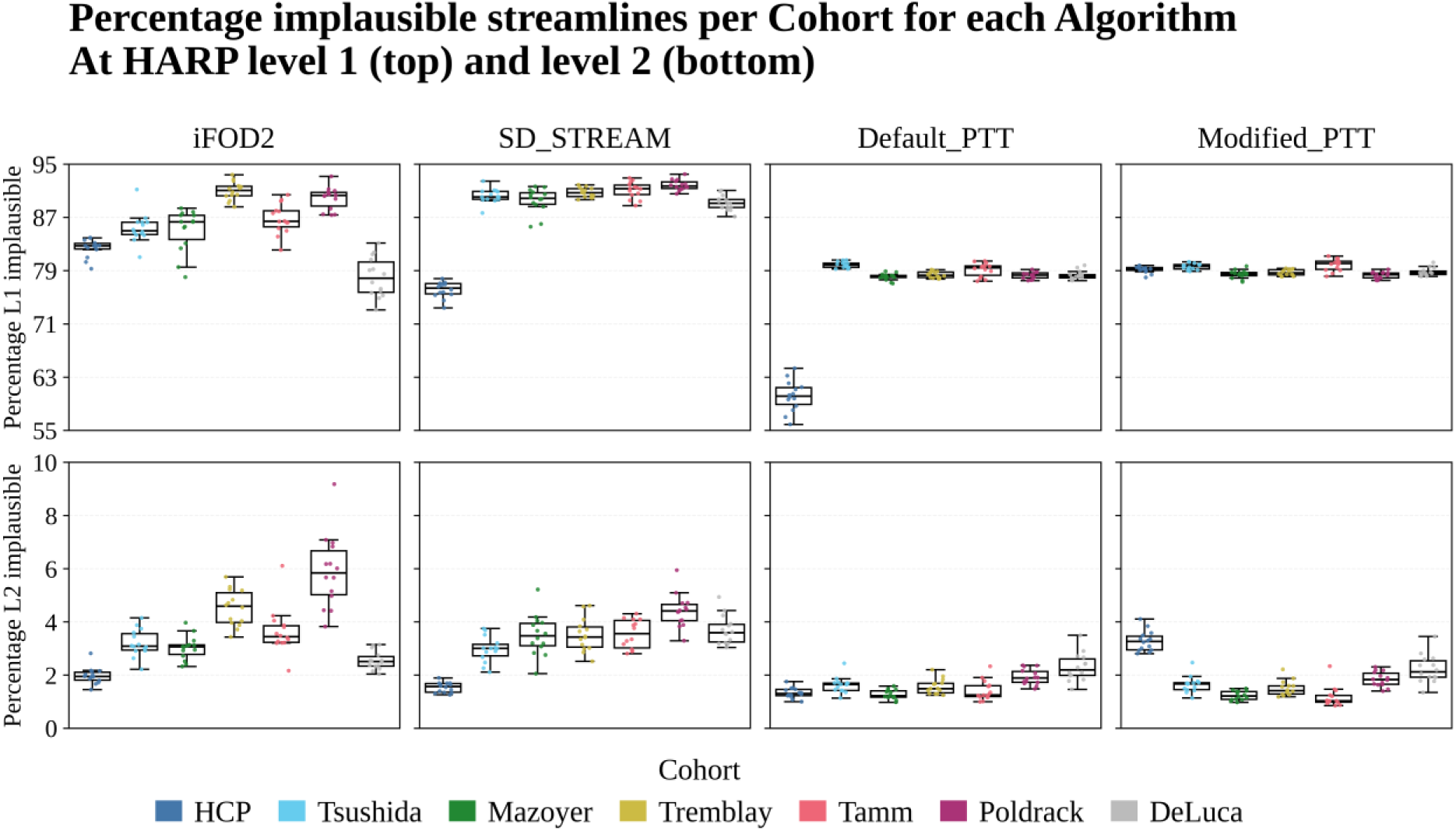
Quantitative ratios of HARP-flagged implausible streamlines. The figure reports, across cohorts and tractography algorithms, the ratio of L1 implausible to L0 plausible streamlines (top panel) and the ratio of L2 implausible to L1 plausible streamlines (bottom panel). Columns correspond to tracking algorithms, and each subplot shows seven cohort-specific boxplots. HARP level 1 rules classify 55.90–93.50% of L0 streamlines as anatomically implausible, with significant main effects of Cohort, Algorithm, and their interaction. Algorithm-specific post-hoc contrasts indicate significant difference between the HCP and all other clinical cohorts within the Default PTT algorithm, but none within the Modified PTT one; more heterogeneous patterns were found for the other algorithms. HARP level 2 rules filter fewer streamlines (0.85–9.18%), and the corresponding statistical model again revealed significant main effects of Cohort, Algorithm, and their interactions. At this level, differences between HCP and clinical cohorts are less pronounced, with fewer significant algorithm-specific post-hoc comparisons.

HARP level 2 rules filter out fewer streamlines than HARP level 1 rules, between 0.85% and 9.18%. The logit-transformed ratio of L2 implausible to L1 plausible streamlines was modeled using a linear mixed-effects model (log-likelihood = 87.51; random-intercept variance = 0.014 ± 0.024). Diagnostic plots indicated good model fit (Supplementary Figure 2). Wald tests identified significant main effects of Cohort, Algorithm, and their interaction. Estimated marginal means further indicated significant differences among all algorithm pairs and among 14/21 cohort pairs. Post-hoc simple-effects analyses showed significant Algorithm differences in 33/42 cohort-specific contrasts and significant Cohort differences in 52/84 algorithm-specific contrasts. In particular, within the iFOD2 algorithm, differences were significant in all pairwise cohort comparisons but for Mazoyer – Tamm, DeLuca – Mazoyer, Mazoyer – Tsushida, Tamm – Tsushida; within the SD_STREAM algorithm, all pairwise post-hoc differences with the HCP cohort were statistically significant, and other comparisons were heterogeneous; within the Default_PTT algorithm, results were mixed; within the Modified_PTT algorithm, all pairwise post-hoc differences with the HCP cohort were statistically significant, and other comparisons were heterogeneous.

The L1 plausible streamlines contain both L2 plausible and L2 implausible streamlines. Figure 6 quantifies the endpoint distribution of both L2 plausible and L2 implausible streamlines with respect to the three L1 endpoint ROIs for the HCP cohort, and all tracking methods. For this cohort, and across algorithms, most plausible streamlines connect two gray matter areas (Top panel), and the large majority of implausible streamline connect two brainstem areas (Middle panel). The type of implausible streamlines found across algorithms is different (Bottom panel). iFOD2 and SD_STREAM include a larger percentage of streamlines matching trj10 (brainstem-to-brainstem connections crossing cerebellar white matter) when compared to both PTT algorithms (Bottom panel, top row). These algorithms also include a larger portion of associations crossing the brainstem (trj 5 and 6) compared to the PTT methods (Bottom panel, middle row). In contrast, the proportion of implausible streamlines connecting two GM areas found with PTT algorithms is more evenly distributed among trj1, trj2, trj5, trj6 and trj7. All algorithms found approximately as many left projections crossing the right hemisphere (trj3) as right projections crossing the left hemisphere (trj4) (Bottom panel, bottom row).

**Figure 6.**
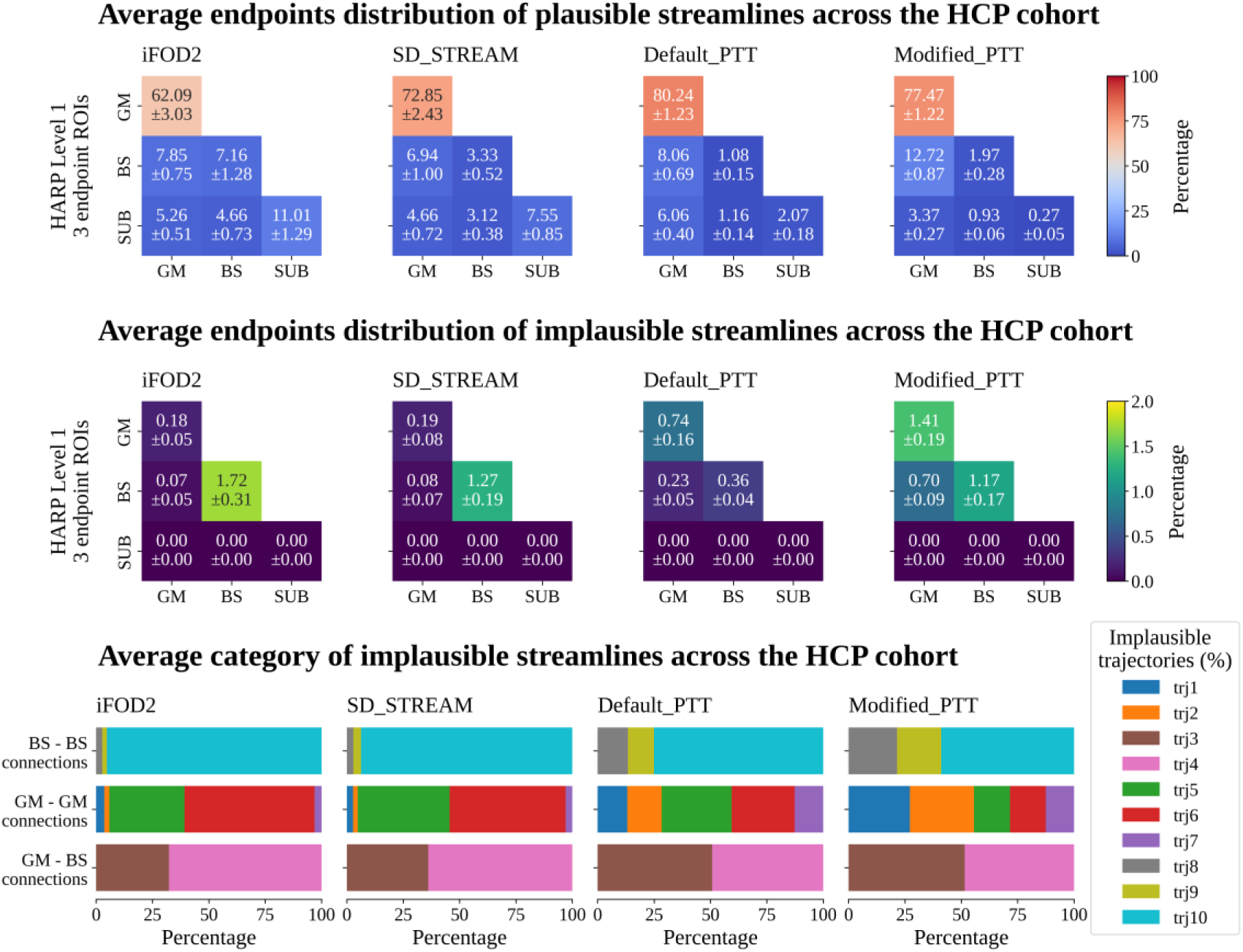
L1 endpoint classification of L2 plausible and implausible streamlines. The figure summarizes, for the HCP cohort and all tracking algorithms, the endpoint distribution of L2 plausible (Top panel) and L2 implausible streamlines (Bottom panel) with respect to the three L1 endpoint ROIs. For each algorithm, the elements of the plausible and implausible endpoint matrices sum to 100. Across methods, plausible streamlines predominantly connect two gray matter regions, whereas most implausible streamlines connect two brainstem regions. The specific types of implausible trajectories (bottom panel) vary by algorithm, with iFOD2 and SD_STREAM showing a higher proportion of brainstem-to-brainstem and brainstem-crossing associations, while PTT algorithms display a more balanced distribution across several GM-based implausible classes. All methods show roughly symmetric proportions of left-to-right and right-to-left misrouted projections.

The 9-elements composition was ilr transformed and modeled using a linear mixed-effects model. Wald tests identified significant main effects of Cohort, Algorithm, and their interaction. Estimated marginal means further indicated significant differences among all algorithm pairs and all cohort pairs. Post-hoc simple-effects analyses showed significant Algorithm differences in 38/42 cohort-specific contrasts and significant Cohort differences in 79/84 algorithm-specific contrasts. In particular, no significant differences between the Default and the Modified PTT algorithms were found for the Mazoyer, Poldrack, Tremblay and Tsushida cohorts.

### 4.3 Comparison with post-hoc filtering techniques

Figure 7 describes the SIFT2 and COMMIT2 weights of the L2 implausible streamlines identified by HARP, across all cohorts for the Default PTT algorithm. The histograms in the top panel show that implausible streamlines can be associated with the full range of harmonized weights for both filtering algorithms. However, there are more implausible streamlines with a large harmonized SIFT2 weight than with a smaller one. Conversely, there are slightly fewer implausible streamlines with a large COMMIT2 harmonized weight than with a smaller one. The histogram of harmonized COMMIT2 weights for the L2 implausible streamlines is overall rather uniform, while the corresponding histogram obtained with SIFT2 weights is uniform only until the harmonized weight of ∼0.95, and rapidly increases after that value.

**Figure 7.**
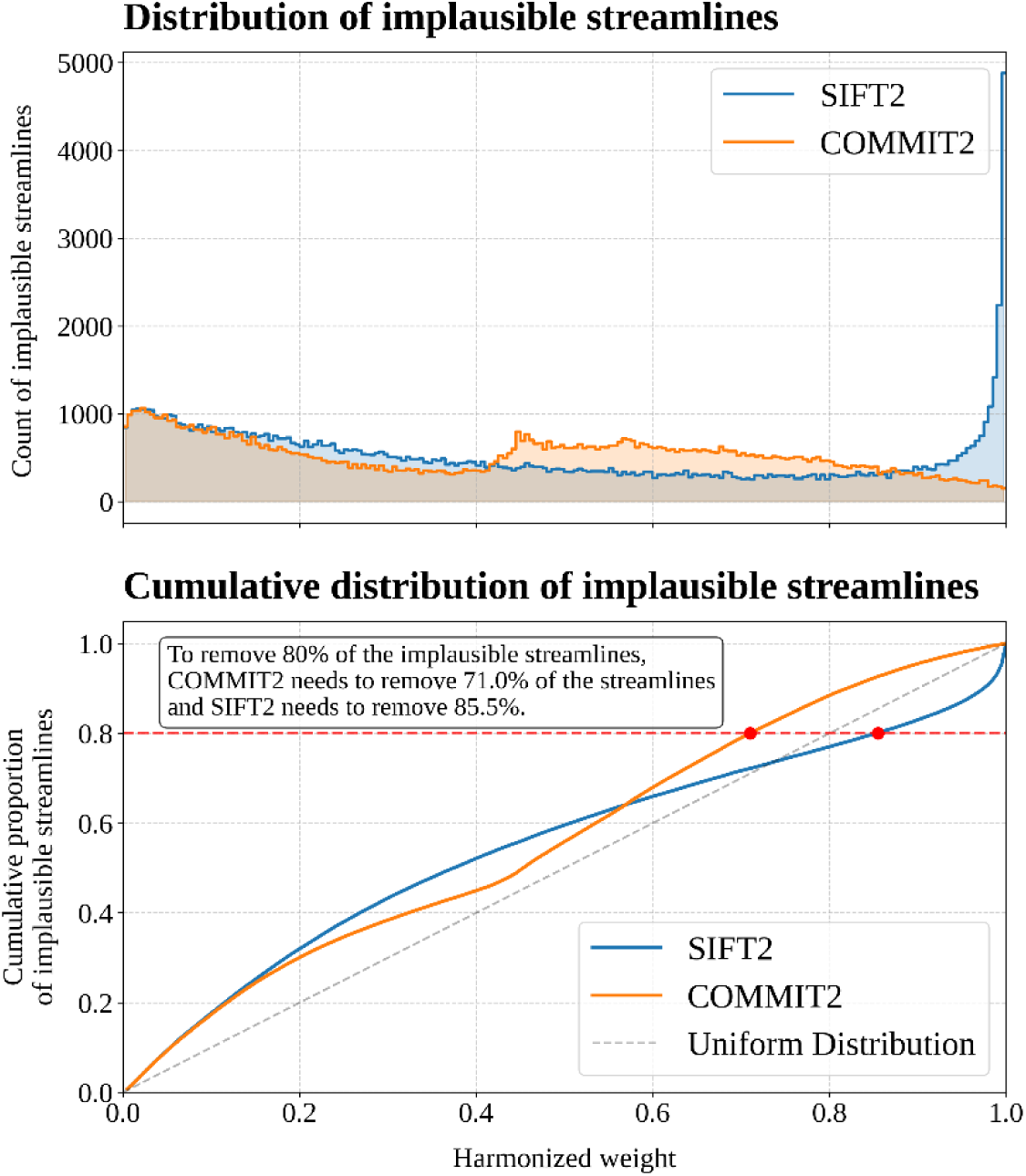
Weights of HARP L2 implausible streamlines under SIFT2 and COMMIT2. The figure summarizes, for all subjects using the Default PTT algorithm, the distribution of harmonized SIFT2 and COMMIT2 weights assigned to L2 implausible streamlines identified by HARP. The top panel histograms show that implausible streamlines span the full range of harmonized weights for both filtering methods, with SIFT2 assigning proportionally larger weights and COMMIT2 showing a more uniform distribution. The bottom panel ECDFs depict the cumulative proportion of implausible streamlines removed as a function of the harmonized weight. The harmonized weight required to capture a given amount of HARP L2 implausible streamlines differs between algorithms.

The bottom panel shows the cumulative proportion of L2 implausible streamlines detected as a function of the harmonized weight. These ECDF curves show that, on average across the 98 subjects, to remove 80% of implausible streamlines, we would need to discard the lowest 71% of streamlines according to the harmonized COMMIT2 weight; and the lowest 85.5% of streamlines according to the harmonized SIFT2 weight.

Figure 8 shows the same analysis performed with stratification over cohort, which produced similar results. Across cohorts, removing 80% of the implausible streamlines consistently required discarding fewer streamlines when ranked by COMMIT2 weights than by SIFT2 weights. Both methods exhibited systematic dependence on acquisition protocol, as reflected in the cohort-stratified ECDFs, which showed an identical top-to-bottom ordering for the two filtering techniques.

**Figure 8.**
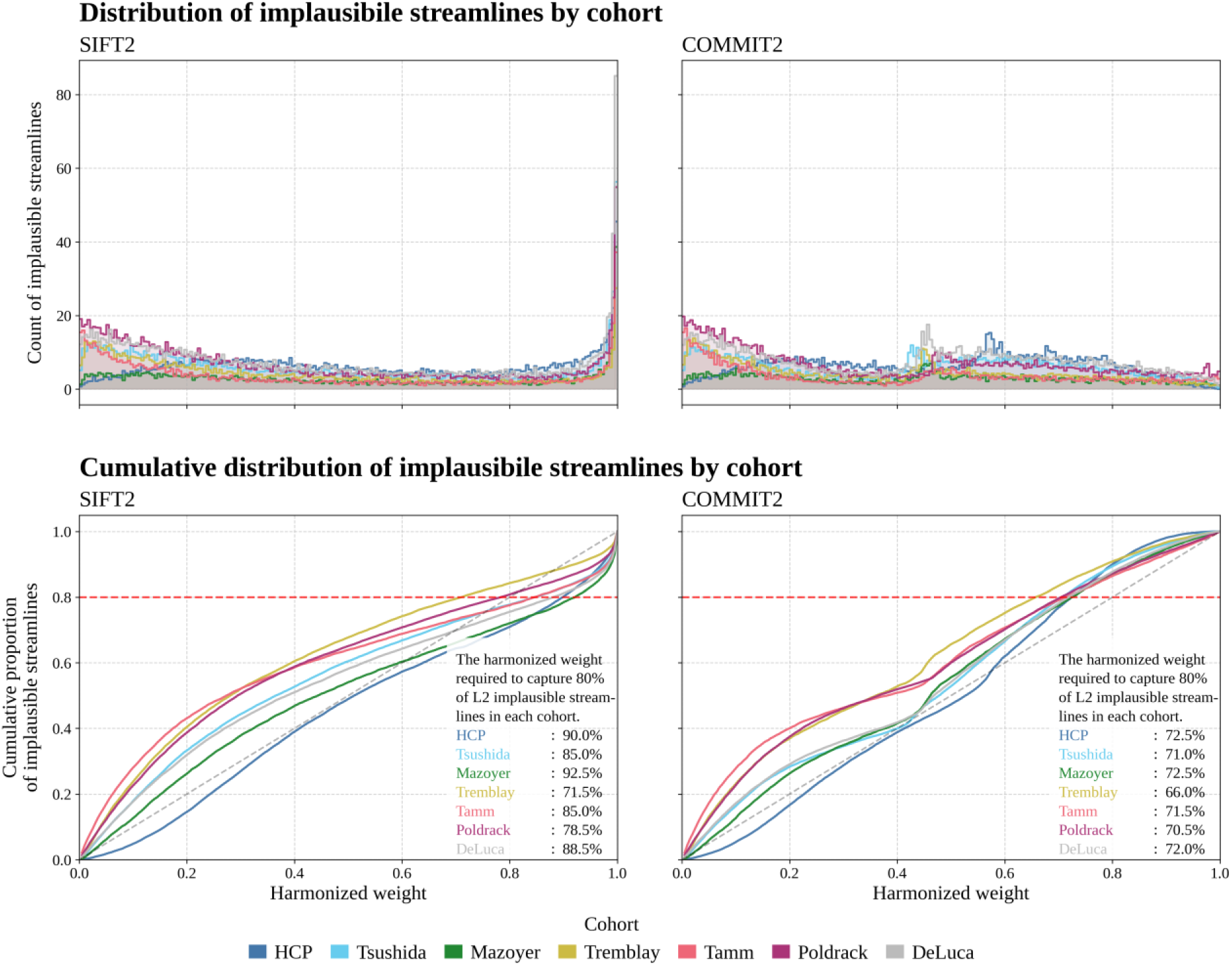
Cohort-stratified weights of HARP L2 implausible streamlines under SIFT2 and COMMIT2. The figure reports, for all subjects and cohorts using the Default PTT algorithm, the cohort-stratified distribution of harmonized SIFT2 and COMMIT2 weights assigned to L2 implausible streamlines identified by HARP. In each panel, curves are color-coded by cohort. As in the pooled analysis, implausible streamlines span the full range of harmonized weights, with SIFT2 assigning proportionally larger weights and COMMIT2 showing a more uniform distribution. The ECDFs in the bottom panel show that, across all cohorts, as an example, removing 80% of L2 implausible streamlines consistently requires discarding fewer streamlines when ranked by COMMIT2 weights than by SIFT2 weights. The ECDF curves related to SIFT2 also appear more different between cohorts. Both filtering methods display a systematic dependence on acquisition protocol, reflected in the consistent top-to-bottom cohort ordering observed across the ECDFs.

## 5. Discussion

We introduce Hierarchical Anatomical Refinement of Pathways (HARP), a rule-based framework that extends Anatomically Constrained Tractography (ACT) (R. E. Smith et al., 2012) by incorporating finer-grained anatomical priors. Our results show that depending on the imaging and tractography pipeline, HARP can reduce false positive streamlines by up to 9%. Furthermore, we also showed that the anatomically implausible streamlines identified by HARP could not be unambiguously distinguished from plausible ones using SIFT2 (R. E. Smith et al., 2015) or COMMIT2 (Schiavi et al., 2020) weights, highlighting the complementarity of post-hoc filtering techniques and anatomically driven constraints.

The comprehensive quantification of the proportion of streamlines deemed implausible (see Fig. 5) was performed on unconstrained (L0) streamlines. Although this approach is nowadays uncommon when studying whole-brain connectivity, it provides two key advantages for the current work: (i) a fair comparison across tractography algorithms that would otherwise internally leverage slightly different anatomical priors and mechanisms; and (ii) a direct quantification of the contribution of ACT rules (HARP level 1) and the additional improvement that comes with the novel HARP level 2 rules. Our systematic evaluation also allowed us to distinguish the significant main effects due to algorithm, cohort, and their interaction at different levels of HARP, which highlights that algorithmic choices and differences in data quality jointly shape the prevalence of implausible trajectories.

### Anatomical basis for implausible HARP level 2 trajectories

HARP level 2 trajectories are considered implausible for three main reasons. First, as we showed in the ISMRM 2015 challenge dataset, whole-brain tractography generated all ten HARP L2 implausible trajectories despite their absence from the curated ground truth bundles. This controlled result demonstrates that such trajectories can arise from the ill-posed nature of tractography itself, rather than from true anatomical connectivity (Jbabdi & Johansen-Berg, 2011; Maier-Hein et al., 2017). Second, these trajectories violate the classical macroscopic organization of white-matter pathways: association fibers are expected to remain within the same hemisphere, commissural fibers cross between hemispheres through forebrain commissural systems rather than the brainstem, and projection fibers are not expected to reach the brainstem by first traversing the contralateral cortical white matter (Catani, 2025; Meynert & Sachs, 1885; Schmahmann & Pandya, 2006). Similarly, brainstem-to-brainstem streamlines making long excursions through cortical or cerebellar white matter are unlikely to represent coherent macroscopic pathways. Third, even if rare anatomical exceptions were eventually described, their existence would not necessarily imply that they are detectable as coherent pathways with dMRI tractography, which operates at a much coarser spatial scale and is known to suffer from both false positive and false negative reconstructions (Aydogan et al., 2018; Maier-Hein et al., 2017; Reveley et al., 2015; Thomas et al., 2014).

Therefore, HARP level 2 implausibility should be interpreted as a conservative rule-based classification: rejected streamlines violate established macroscopic anatomical priors at the current segmentation level, whereas retained streamlines are not guaranteed to be biologically valid.

### Effect of HARP’s level

Our results indicate that HARP level 1 (equivalent to ACT) was the most impactful step, flagging as implausible between 55.90% and 93.50% of all L0 streamlines across cohorts and algorithms (see Fig. 5, top row). This magnitude is expected, as level 1 rules address the most frequent errors occurring when performing anatomically unconstrained tractography tracking, even in high-quality data. These errors primarily include premature termination in WM (Girard et al., 2014; Jeurissen et al., 2019; R. E. Smith et al., 2012), and spurious propagation into CSF. The former problem is prominent (Supplementary Fig. 3) owing to the complexity of white matter organization, for example, due to the high prevalence of crossing fibers in WM, which is reported to be present in up to 90% of voxels (Jeurissen et al., 2013).

Among the L1 plausible streamlines, L2 rules flagged between 0.85–9.18% of streamlines as implausible (see Fig. 5, bottom row). This value is smaller compared to the percentage of streamlines that L1 rules removed from an unconstrained tractogram. The modest additional pruning by L2 is expected given the finer-grained definition of rules, based on both endpoints and traversed regions (see Fig. 2). Consequently, we expect HARP to remove fewer streamlines at higher levels.

While assessing the impact of the seeding strategy on the proportion of L2 implausible streamlines was outside the scope of this work, it is important to consider this factor as a source of variability. We seeded in the white matter rather than at the GM-WM interface to ensure whole brain coverage (Jeurissen et al., 2019). This choice is known to promote long streamlines (Girard et al., 2014; R. E. Smith et al., 2012), potentially increasing the prevalence of implausible trajectories described in Figure 2.

### Effect of tractography algorithm on HARP

Estimated marginal means and post-hoc contrasts demonstrated systematic differences among all algorithms at both HARP levels of L1 and L2. Two patterns are particularly informative:

- Both PTT-based approaches produced fewer implausible streamlines and showed lower variability than iFOD2 and SD_STREAM (see Fig. 5). This mirrors prior benchmarking (Aydogan & Shi, 2021; Girard et al., 2023) showing that PTT’s built-in topographic and geometric regularization preserves bundle coherence and discourages sharp deviations, helping to yield streamlines that are more compliant with HARP rules.
- Deterministic SD_STREAM produced more level 1 implausible streamlines than probabilistic iFOD2, which can be attributed to probabilistic methods’ ability to consider alternative plausible orientations in complex WM areas (Descoteaux et al., 2009). Deterministic tracking, by following an optimal-direction approach at each step, may be more vulnerable to errors in crossing regions, increasing L1 implausible streamlines (Descoteaux et al., 2009). At level 2, however, the proportion of implausible streamlines was higher for iFOD2 than for SD_STREAM, suggesting that deterministic tracking may be more robust to the specific endpoint- and pathway-based constraints enforced at level 2.

### Effect of cohort on HARP

Cohort effects were pronounced. Post-hoc simple-effects analyses showed that the HCP cohort differed significantly from all clinical cohorts at HARP level 1 across all algorithms except Modified PTT; and at level 2 across all algorithms except Default PTT. These differences are consistent with the higher quality of HCP acquisitions (Van Essen et al., 2012). However, the cohort effect depended on algorithm (see Fig. 5):

- The strong HCP–clinical differences at HARP level 1 occurring for the Default PTT algorithm did not occur when using Modified PTT algorithm, suggesting that parameter adjustments can partly compensate for lower quality or heterogeneous clinical protocols. On the other hand, the result may suggest that the Default PTT settings are preferable for the HCP data compared to the Modified PTT parameters.
- A larger variability across cohorts was observed for MRtrix3 algorithms (iFOD2 and SD_STREAM) than for PTT ones. PTT priors may mitigate the variability present in noisier, lower-resolution clinical data, narrowing cohort differences.

### Endpoint classification differences

Our endpoint classification analysis reveals clear, algorithm-dependent differences in how L2 plausible and implausible streamlines are classified in the HCP cohort (see Fig. 6). The compositional analysis across all datasets confirms that these differences are systematic: both cohort and algorithm exert significant main effects, with a significant interaction, and nearly all cohort-specific and algorithm-specific contrasts are significant.

Two mechanisms likely give rise to these different endpoint classifications. First, L1 rules may exert a disproportionate impact on a specific subset of trajectories. This is consistent with the different amount of L1 implausible streamlines observed in Figure 5 and indicates that L1 constraints may shape downstream L2 streamlines endpoint classification. Second, algorithm-specific tracking parameters influence streamline course and termination, thereby biasing which classification streamlines will ultimately have. Because endpoint ROIs and rules were held constant across algorithms, these differences are best attributed to the chosen tractography algorithm.

Within the HCP cohort, the observed predominance of L2 plausible GM-to-GM connections across algorithms (see Fig. 6, bottom row) is expected as this ROI is much larger than other ROIs. Conversely, the concentration of L2 implausible streamlines in the BS-BS class follows from two features of our framework: several HARP L2 rules specifically target brainstem-related connections; and the brainstem is the ROI most susceptible to ambiguous segmentation given its shape, location and proximity to other ROIs, even with high-quality imaging data (Sclocco et al., 2018; Tang et al., 2018).

### Characterization of HARP implausible streamlines through post-hoc filtering algorithms

Comparisons between SIFT2 and COMMIT2 have previously been reported in several studies, primarily focusing on their differential effects on connectomics metrics rather than on individual streamline weights (Frigo et al., 2020; He et al., 2024; Koch et al., 2022; Sarwar et al., 2023; Wang et al., 2025). In contrast, our study allows for a clear and straightforward comparison of SIFT2 and COMMIT2 at the single-streamline level, through the harmonized weights shown in figures 7-8. The harmonized weights simply sort the streamlines in terms of their rank obtained with these post-hoc filtering methods. Notably, we observe that HARP-identified L2 implausible streamlines span the full range of harmonized weights under both SIFT2 and COMMIT2 (figures 7-8).

These findings have two immediate implications. First, low harmonized weights (ranks) do not directly signal anatomical implausibility, and no clear-cut harmonized weight threshold could separate L2 plausible from L2 implausible streamlines. Moreover, even some HARP L2 plausible streamlines receive low harmonized COMMIT2 or SIFT2 weights, which can be partly explained by seeding strategy since the whole WM seeding biases the tractogram toward longer trajectories (Girard et al., 2014; R. E. Smith et al., 2012) that tend to receive lower post-hoc weights. Thus, quantitative weights and anatomical plausibility labels are related but not interchangeable.

Second, there remains a clear role for SIFT2, COMMIT2, and related approaches to obtain quantitative insights. These methods inherently aim for global objectives, in that the weight assigned to any given streamline depends on the presence of all other streamlines. HARP does not provide quantitative information but can substantially constrain the input tractogram on anatomical grounds. Our results therefore support a workflow in which HARP is applied first to exclude anatomically implausible streamlines, followed by post-hoc filtering to derive quantitative measures from the anatomically curated set.

Importantly, our results do not identify a universally better post-hoc filtering strategy. The two methods optimize different functions, and the appropriate choice depends on the research question. At the same time, COMMIT2 appears more aligned with our anatomical rules in this context as observed in figures 7-8.

### HARP’s rules: Relationship to prior work

The works most closely related to HARP are ACT (R. E. Smith et al., 2012) and ExTractorFlow (Petit et al., 2019, 2023), as both introduce a set of anatomical priors to constrain tractography. As confirmed by our results (see Fig. 5), the adoption of ACT rules is highly effective to ensure the anatomical plausibility of whole-brain tractograms across cohorts and tracking algorithms. However, because we started with unconstrained tractograms (L0), HARP level 1 implementation in the current work differs from ACT in two specific aspects.

First, in ACT, streamlines are not allowed to traverse subcortical gray matter and are forcibly terminated before exiting these regions. While appropriate in some cases, this constraint is not aligned with many studies explicitly focusing on subcortical connectivity, where streamlines are required to fully traverse subcortical structures (Behrens, Johansen-Berg, et al., 2003; Draganski et al., 2008; Palesi et al., 2015). HARP instead allows either termination within subcortical regions or complete crossing through these regions if the FOD supports such trajectories. This modification, already proposed in (Bajada et al., 2019), may help avoid implicitly inflating the number of subcortical connections by splitting streamlines at their boundaries. Second, ACT truncates streamlines immediately upon entering cortical gray matter. As already noted in the original ACT paper, there is no anatomical reason to prevent streamlines from projecting into this region. At the same time, this is difficult to implement using voxel-based anatomical priors. By including surface-based priors in our whole-brain tractography pipeline, we were able to track through the cortical gray matter already at level 0.

Similar to ACT, HARP allows for flexible handling of abnormal or pathological tissue and can in principle be adopted in global tractography algorithms. More generally, ACT rules are not exhaustive and can be complemented, as demonstrated in (Petit et al., 2019, 2023). Petit et al., 2019, first introduced ExTractorFlow, a hierarchical filtering framework and evaluated its impact across multiple tracking algorithms and two datasets (HCP and Mazoyer), both included in the current study as well. Petit et al., 2023, builds on the Petit et al., 2019 framework and applies expanded rules to the full Mazoyer cohort but does not include most of HARP’s constraints (rules 5–10) and it is also limited to the analysis of the angular gyrus. It does, however, add rules for gyral stem crossings, Boolean plausibility criteria based on Johns Hopkins University template ROIs, along with manually segmented ROIs specific to the angular gyrus.

### HARP: need & scope

By definition, dMRI-based tractography is an ill-posed problem: given only local diffusion measurements, the space of possible long-range connections is too vast, and multiple fiber configurations are equally plausible (Daducci et al., 2025). As a result, tractograms that rely on minimal constraints are intrinsically prone to errors (Thomas et al., 2014). Thus, additional constraints are needed to reduce the solution space and guide reconstruction toward anatomically meaningful connections. In particular, the incorporation of anatomical constraints derived from external *a priori* knowledge has consistently been shown to improve tractography performance. Among the tractography variables systematically evaluated by (Aydogan et al., 2018), the use of anatomical constraints emerged as the most influential factor affecting the accuracy of the resulting tractograms. Similar trends were observed in the IronTract challenge, where the post-hoc application of anatomically defined ROIs led to robust reconstruction of pathways across both training and validation injection sites (Maffei et al., 2022). Importantly, (K. G. Schilling et al., 2020) demonstrated that, as increasingly informative priors are introduced into bundle definitions, tractography results converge more closely toward neuronal tracing findings. Although such studies do not target human whole-brain tractography, they provide compelling evidence that carefully designed anatomical constraints can substantially enhance tractography specificity.

It would be highly valuable to extend the use of anatomical priors to the whole brain setting with strategies that can integrate and iteratively refine constraints designed for specific brain structures. Fixed rule-based approaches, such as ACT and ExTractorFlow, provide well-organized frameworks, but they are limited in their ability to adapt as additional or more detailed priors become available. In this context, HARP fills an important gap by providing a scalable solution: its structured yet extendable refinement scheme is specifically designed to iteratively constrain the tractograms while remaining compatible with diverse sources of anatomical information. HARP is therefore not a replacement for existing anatomical constraint-based methods, but rather a general strategy for systematically strengthening and harmonizing anatomical constraints at the whole-brain level.

The primary scope of HARP is improving tractogram specificity. By construction, HARP focuses on identifying and removing false-positive streamlines, which violate anatomical plausibility or consistency based on prior knowledge. Because HARP does not encode explicit rules for detecting missing connections, it does not directly address the widespread issue of false negatives connections (Aydogan et al., 2018). As a result, HARP does not provide a direct mechanism for increasing sensitivity. However, sensitivity and specificity are intrinsically coupled, and improvements in one metric indirectly influence the other (K. G. Schilling et al., 2020).

When false positives can be systematically filtered using HARP, initial tractography can be performed with more permissive parameters, allowing a broader sampling of potential connections across the brain. This reduces the risk of excluding valid pathways during streamline generation, decreasing false negatives. Within this workflow, HARP’s rules flag and remove anatomical implausible connections, thereby shifting the sensitivity–specificity trade-off (Zalesky et al., 2016) toward higher overall sensitivity without sacrificing specificity. Nevertheless, this benefit is indirect: HARP does not offer a dedicated strategy for identifying or correcting false negatives, as it does for false positives.

### Limitations

A limitation of the proposed HARP framework lies in the necessary simplifications underlying the definition of the anatomical rules given a set of available ROIs. While designed to be intuitive and broadly applicable, these rules are likely not exhaustive, which may lead to streamline misclassification. For instance, no dedicated rules were defined for subcortical regions.

Second, the performance of HARP critically depends on the quality and specificity of the segmented ROIs. Whether using surface meshes, binary image masks or partial volume fractions, the rules are identical, and our software implementation supports all these options. However, each segmentation method comes with specific drawbacks. While we adopted high-resolution surface meshes to limit partial volume effects and discretization errors arising with image-based approaches, such choice is prone to intersecting regions where ROIs were no longer mutually exclusive. In our implementation, to circumvent these ambiguities, we used a priority order to uniquely classify each streamline into only one category. In this study, we observed that on average 2.71% of streamlines classified as plausible at level 2 were ambiguous due to overlaps in segmented ROIs, highlighting the sensitivity of the approach to ROI definition. While this percentage remains relatively small, it nevertheless indicates that segmentation uncertainty influences the outcome and should be checked during quality control.

### Future directions

First, HARP can be readily adopted by the community to further refine already curated tractograms, providing an additional layer of anatomical plausibility assessment that is transparent and easy to interpret.

Second, as an alternative to post-hoc filtering, HARP could be implemented for use during fiber tracking. Applying anatomical plausibility constraints during streamline generation reduces computational efficiency.

Third, the current rule set represents an initial and limited formalization of anatomical knowledge. Future work should focus on translating additional well-established anatomical observations and principles into further HARP rules. For instance, the presence of heterotopic callosal connections is well-established: shown in the macaque’s visual callosal connections (e.g., Van Essen et al., 1982) and then extended to other areas (Rizzo et al., 2026), heterotopic callosal connections have also been reconstructed using tractography in humans (Szczupak et al., 2023). Thus, callosal fibers could be further categorized into homo- and heterotopic connections based on their geometry (De Benedictis et al., 2016; Innocenti et al., 2022; Lent et al., 2026). A further category of crossed cortico-striatal projections could help mirror results found in macaque (Borra et al., 2022; Innocenti et al., 2017). Improved segmentation of the cerebellum, brainstem, and associated nuclei could also draw on and apply rules targeting well-known pathways involving the deep cerebellar nuclei, superior cerebellar peduncle, olivocerebellar projections, and auditory pathways (Basile et al., 2020; Krimmel et al., 2025; Li et al., 2024). Moreover, some brain regions in these areas exhibit reciprocal connectivity, whereas others show predominantly unidirectional projections (Li et al., 2024). Leveraging this neuroanatomical information could add directionality to the connections at higher levels of HARP, a distinct improvement given that tractography inherently lacks directionality (Jbabdi & Johansen-Berg, 2011).

Other anatomical insights remain challenging to encode within tractography frameworks. Translating detailed anatomical knowledge into tractography priors is in fact one of the most pressing challenges for the field as noted in the Millennium Pathways for Tractography (Descoteaux et al., 2025). Beyond the difficulty of mapping focal tracer injection sites onto comparatively large in vivo MRI voxels, many connectional properties described in invasive animal studies, such as complex divergence of layer-specific projections (Coudé et al., 2018; Kita & Kita, 2012; Parent & Parent, 2005), do not readily transfer to tractography, due to the fundamental mismatch between data from single-neurons/axon measurements and the fiber bundles represented by streamlines (Jones et al., 2013). To keep in mind as well, neuroanatomy is actively evolving with new methods (e.g., auf der Heiden et al., 2025; Cooper et al., 2026), which can be expected to influence continued developments in tractography.

## 6. Conclusions

HARP provides an extensible framework for incorporating prior anatomical knowledge in a manner that can be iteratively refined to reduce false positives. Its demonstrated effectiveness, acknowledged limitations, and dependence on anatomical priors highlight the need for collective evaluation and continued development, rather than treating the specific rules proposed in the current work as definitive. We therefore view HARP not as a closed or finalized methodology, but as an open invitation to the tractography and wider community: to scrutinize existing rules, propose new ones, and collaboratively advance anatomically informed tractography. Through shared efforts, such principled constraints have the potential to substantially improve the biological interpretability and reliability of tractography-based analyses.

## 7. Data and code availability

Data used in this paper were provided in part by the Human Connectome Project, WU-Minn Consortium (Principal Investigators: David Van Essen and Kamil Ugurbil; 1U54MH091657) funded by the 16 NIH Institutes and Centers that support the NIH Blueprint for Neuroscience Research; and by the McDonnell Center for Systems Neuroscience at Washington University.

The TractoInferno and the ISMRM 2015 datasets used in this study are publicly available from the original sources cited in the manuscript. The software implementing HARP is publicly available in Trekker’s harp GitHub branch (https://github.com/dmritrekker/trekker/tree/harp).

## 9.#Acknowledgements

SL received funding from the Instrumentariumin Tiedesäätiö sr. DBA received funding from the Research Council of Finland (grant numbers 348631 and 353798) and the Technology Industries of Finland Centennial Foundation (grant number 4302). The computations presented above were performed using computing resources within the Aalto University School of Science “Science-IT” project. Supplementary Figure 3 was made through SankeyMATIC.com.

## 10. CRediT authorship contribution statement

**Simona Leserri:** Data curation, Formal analysis, Funding acquisition, Investigation, Methodology, Software, Validation, Visualization, Writing – original draft, Writing – review and editing. **Kathleen S. Rockland:** Investigation, Validation, Writing – review and editing. **Dogu Baran Aydogan:** Conceptualization, Data curation, Formal analysis, Funding acquisition, Investigation, Methodology, Project administration, Resources, Software, Supervision, Validation, Visualization, Writing –original draft, Writing – review and editing.

## 11. AI disclosure

ChatGPT-5.1 and Copilot ChatGPT-5.3 were used during the preparation of this article. All AI-generated suggestions were carefully reviewed and edited by the authors. The authors take full responsibility for the content of the publication.

## Appendix

### Criteria

1. Streamline enters CSF
2. Streamline with both ends in any L1 ROIs
3. Streamline enters LN_WM
4. Streamline enters RN_WM
5. Streamline enters CER_WM
6. Streamline enters BS
7. Streamline has one endpoint in LN_GM and the other endpoint in LN_GM
8. Streamline has one endpoint in LN_GM and the other endpoint in RN_GM
9. Streamline has one endpoint in LN_GM and the other endpoint in L_SUB
10. Streamline has one endpoint in LN_GM and the other endpoint in R_SUB
11. Streamline has one endpoint in LN_GM and the other endpoint in CER_GM
12. Streamline has one endpoint in LN_GM and the other endpoint in BS
13. Streamline has one endpoint in RN_GM and the other endpoint in RN_GM
14. Streamline has one endpoint in RN_GM and the other endpoint in L_SUB
15. Streamline has one endpoint in RN_GM and the other endpoint in R_SUB
16. Streamline has one endpoint in RN_GM and the other endpoint in CER_GM
17. Streamline has one endpoint in RN_GM and the other endpoint in BS
18. Streamline has one endpoint in L_SUB and the other endpoint in L_SUB
19. Streamline has one endpoint in L_SUB and the other endpoint in R_SUB
20. Streamline has one endpoint in L_SUB and the other endpoint in CER_GM
21. Streamline has one endpoint in L_SUB and the other endpoint in BS
22. Streamline has one endpoint in R_SUB and the other endpoint in R_SUB
23. Streamline has one endpoint in R_SUB and the other endpoint in CER_GM
24. Streamline has one endpoint in R_SUB and the other endpoint in BS
25. Streamline has one endpoint in CER_GM and the other endpoint in CER_GM
26. Streamline has one endpoint in CER_GM and the other endpoint in BS
27. Streamline has one endpoint in BS and the other endpoint in BS

### Priority order

1. One endpoint in LN_GM and the other endpoint in LN_GM
2. One endpoint in LN_GM and the other endpoint in RN_GM
3. One endpoint in LN_GM and the other endpoint in L_SUB
4. One endpoint in LN_GM and the other endpoint in R_SUB
5. One endpoint in LN_GM and the other endpoint in CER_GM
6. One endpoint in LN_GM and the other endpoint in BS
7. One endpoint in RN_GM and the other endpoint in RN_GM
8. One endpoint in RN_GM and the other endpoint in L_SUB
9. One endpoint in RN_GM and the other endpoint in R_SUB
10. One endpoint in RN_GM and the other endpoint in CER_GM
11. One endpoint in RN_GM and the other endpoint in BS
12. One endpoint in L_SUB and the other endpoint in L_SUB
13. One endpoint in L_SUB and the other endpoint in R_SUB
14. One endpoint in L_SUB and the other endpoint in CER_GM
15. One endpoint in L_SUB and the other endpoint in BS
16. One endpoint in R_SUB and the other endpoint in R_SUB
17. One endpoint in R_SUB and the other endpoint in CER_GM
18. One endpoint in R_SUB and the other endpoint in BS
19. One endpoint in CER_GM and the other endpoint in CER_GM
20. One endpoint in CER_GM and the other endpoint in BS
21. One endpoint in BS and the other endpoint in BS

## Supplementary

**Supplementary Figure 1:**
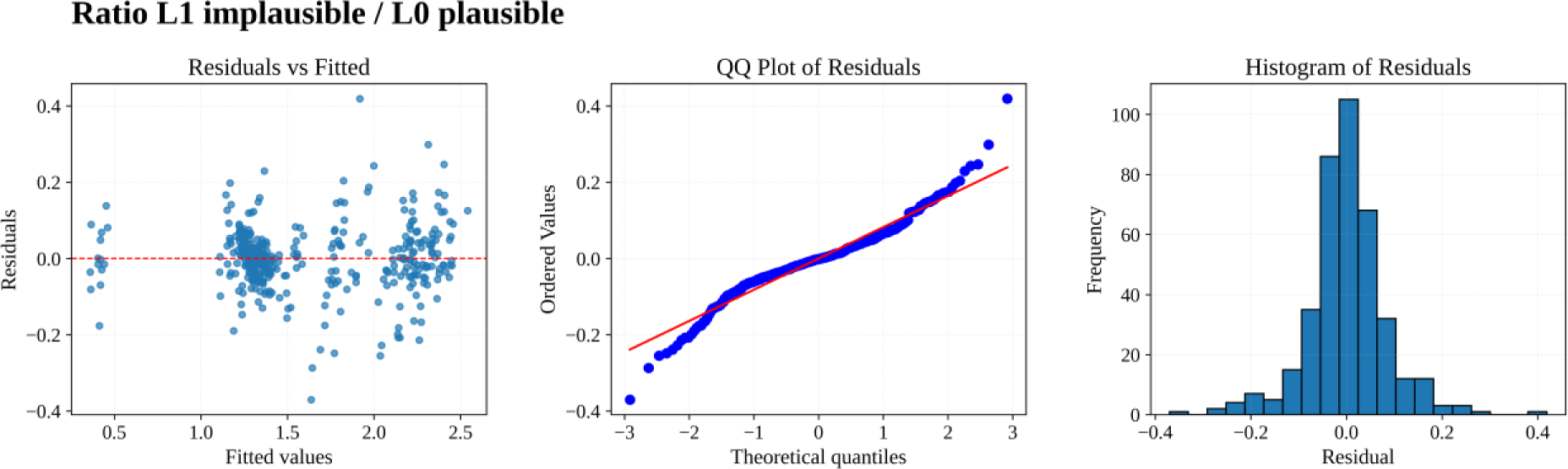
Diagnostic plots for the linear mixed model fitted to the logit transformed ratio of L1 implausible/L0 plausible. These plots enable a visual assessment of model fit and underlying assumptions.

**Supplementary Figure 2:**
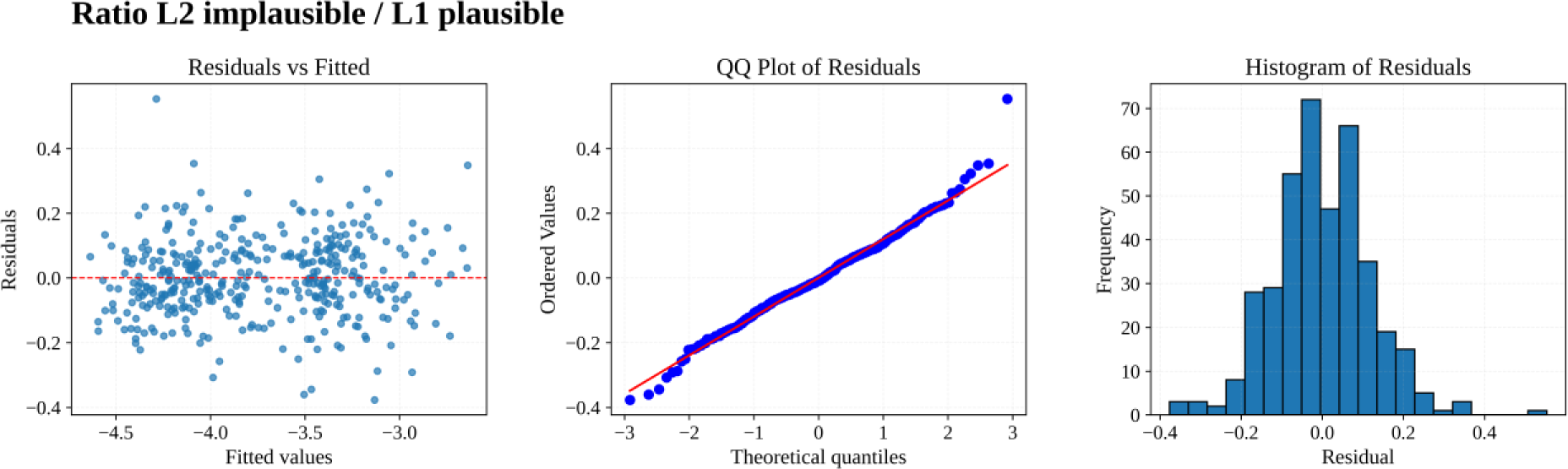
Diagnostic plots for the linear mixed model fitted to the logit transformed ratio of L2 implausible/L1 plausible. These plots enable a visual assessment of model fit and underlying assumptions.

**Supplementary Figure 3.**
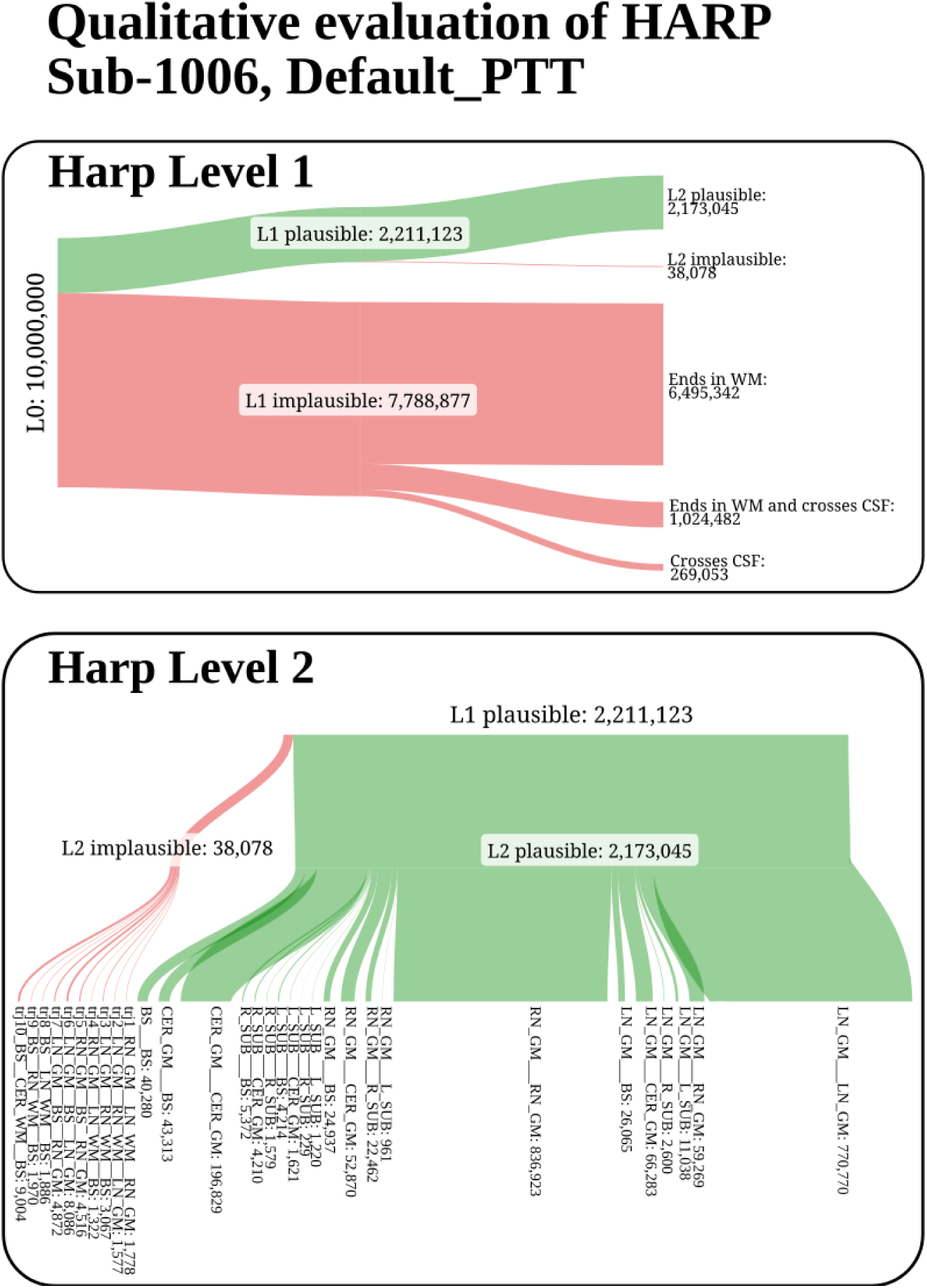
Quantitative effect of HARP on whole-brain tractography. The figure illustrates the impact of applying HARP rules to the tractogram of a representative TractoInferno subject obtained with the Default PTT algorithm. The large majority of HARP level 1 implausible streamlines end in the WM. The ratio of implausible streamlines at level 1 (L1 implausible/L0 %) is larger that the ratio of implausible streamlines at level 2 (L2 implausible/L1 plausible %).

## 12.#References

1. Andersson, J. L. R., Skare, S., & Ashburner, J. (2003). How to correct susceptibility distortions in spin-echo echo-planar images: application to diffusion tensor imaging. NeuroImage, 20(2), 870–888. 10.1016/S1053-8119(03)00336-7

2. Andersson, J. L. R., & Sotiropoulos, S. N. (2016). An integrated approach to correction for off-resonance effects and subject movement in diffusion MR imaging. NeuroImage, 125, 1063–1078. 10.1016/J.NEUROIMAGE.2015.10.019

3. auf der Heiden, F., Axer, M., Amunts, K., & Menzel, M. (2025). Scattering polarimetry enables correlative nerve fiber imaging and multimodal analysis. Scientific Reports 2025 15:1, 15(1), 18493-. 10.1038/s41598-025-02762-w

4. Aydogan, D. B., Jacobs, R., Dulawa, S., Thompson, S. L., Francois, M. C., Toga, A. W., Dong, H., Knowles, J. A., & Shi, Y. (2018). When tractography meets tracer injections: a systematic study of trends and variation sources of diffusion-based connectivity. Brain Structure and Function, 223(6), 2841–2858. 10.1007/S00429-018-1663-8

5. Aydogan, D. B., & Shi, Y. (2021). Parallel Transport Tractography. IEEE Transactions on Medical Imaging, 40(2), 635–647. 10.1109/TMI.2020.3034038

6. Bajada, C., Da, L., Campos, C., Schreiber, J., Muscat, R., & Caspers, S. (2019). A novel constraint for anatomical tractography in the brainstem. *In:* 25th Conference of the Organization for Human Brain Mapping, Roma, 2019. https://www.um.edu.mt/library/oar/handle/123456789/59840

7. Basile, G. A., Quartu, M., Bertino, S., Serra, M. P., Boi, M., Bramanti, A., Anastasi, G. P., Milardi, D., & Cacciola, A. (2020). Red nucleus structure and function: from anatomy to clinical neurosciences. Brain Structure and Function 2020 226:1, 226(1), 69–91. 10.1007/S00429-020-02171-X

8. Behrens, T. E. J., Johansen-Berg, H., Woolrich, M. W., Smith, S. M., Wheeler-Kingshott, C. A. M., Boulby, P. A., Barker, G. J., Sillery, E. L., Sheehan, K., Ciccarelli, O., Thompson, A. J., Brady, J. M., & Matthews, P. M. (2003). Non-invasive mapping of connections between human thalamus and cortex using diffusion imaging. Nature Neuroscience 2003 6:7, 6(7), 750–757. 10.1038/nn1075

9. Behrens, T. E. J., Woolrich, M. W., Jenkinson, M., Johansen-Berg, H., Nunes, R. G., Clare, S., Matthews, P. M., Brady, J. M., & Smith, S. M. (2003). Characterization and propagation of uncertainty in diffusion-weighted MR imaging. Magnetic Resonance in Medicine, 50(5), 1077–1088. 10.1002/MRM.10609

10. Borra, E., Biancheri, D., Rizzo, M., Leonardi, F., & Luppino, G. (2022). Crossed Corticostriatal Projections in the Macaque Brain. Journal of Neuroscience, 42(37), 7060–7076. 10.1523/JNEUROSCI.0071-22.2022

11. Bullmore, E., & Sporns, O. (2012). The economy of brain network organization. Nature Reviews Neuroscience 2012 13:5, 13(5), 336–349. 10.1038/nrn3214

12. Calabrese, E. (2016). Diffusion tractography in deep brain stimulation surgery: A review. Frontiers in Neuroanatomy, 10(MAY), 184199. 10.3389/FNANA.2016.00045/BIBTEX

13. Catani, M. (2025). The brain and its pathways. *Handbook of Diffusion MR Tractography: Imaging Methods, Biophysical Models*, Algorithms and Applications, 3–18. 10.1016/B978-0-12-818894-1.00034-3

14. Ciccarelli, O., Catani, M., Johansen-Berg, H., Clark, C., & Thompson, A. (2008). Diffusion-based tractography in neurological disorders: concepts, applications, and future developments. The Lancet Neurology, 7(8), 715–727. 10.1016/S1474-4422(08)70163-7

15. Cooper, M. L., Selles, M. C., Cammer, M., Redd, C., Gildea, H. K., Sall, J., Chiurri, K. E., Cheung, P., Wheeler, D. G., Saab, A. S., Liddelow, S. A., & Chao, M. V. (2026). Astrocytes connect specific brain regions through plastic networks. Nature 2026, 1–9. 10.1038/s41586-026-10426-6

16. Coudé, D., Parent, A., & Parent, M. (2018). Single-axon tracing of the corticosubthalamic hyperdirect pathway in primates. Brain Structure and Function, 223(9), 3959–3973. 10.1007/S00429-018-1726-X/TABLES/2

17. Daducci, A., Schiavi, S., Christiaens, D., Smith, R., & Alexander, D. C. (2025). Global tractography. *Handbook of Diffusion MR Tractography: Imaging Methods, Biophysical Models*, Algorithms and Applications, 297–314. 10.1016/B978-0-12-818894-1.00014-8

18. De Benedictis, A., Petit, L., Descoteaux, M., Marras, C. E., Barbareschi, M., Corsini, F., Dallabona, M., Chioffi, F., & Sarubbo, S. (2016). New insights in the homotopic and heterotopic connectivity of the frontal portion of the human corpus callosum revealed by microdissection and diffusion tractography. Human Brain Mapping, 37(12), 4718–4735. 10.1002/HBM.23339

19. Dell’Acqua, F., Leemans, A., Zhang, F., Rheault, F., Farquharson, S., & De Luca, A. (2025). The tractographer’s dilemma: understanding sources of variability in tractography. Brain Structure and Function, 230(6), 99-. 10.1007/S00429-025-02964-Y/FIGURES/3

20. Descoteaux, M., Deriche, R., Knösche, T. R., & Anwander, A. (2009). Deterministic and probabilistic tractography based on complex fibre orientation distributions. IEEE Transactions on Medical Imaging, 28(2), 269–286. 10.1109/TMI.2008.2004424

21. Descoteaux, M., Schilling, K. G., Baran Aydogan, D., Beaulieu, C., Borra, E., Chamberland, M., Daducci, A., De Luca, A., Dell’Acqua, F., Dubois, J., Dyrby, T. B., Farquharson, S., Forkel, S., Froeling, M., Griffa, A., Grotheer, M., Guevara, P., Haber, S. N., Kumar Jangir, V., … Petit, L. (2025). Millennium Pathways for Tractography: 40 grand challenges to shape the future of tractography. ArXiv, 10, 23. https://tractography.io/tractanat_retreat/

22. Desikan, R. S., Ségonne, F., Fischl, B., Quinn, B. T., Dickerson, B. C., Blacker, D., Buckner, R. L., Dale, A. M., Maguire, R. P., Hyman, B. T., Albert, M. S., & Killiany, R. J. (2006). An automated labeling system for subdividing the human cerebral cortex on MRI scans into gyral based regions of interest. NeuroImage, 31(3), 968–980. 10.1016/J.NEUROIMAGE.2006.01.021

23. Draganski, B., Kherif, F., Klöppel, S., Cook, P. A., Alexander, D. C., Parker, G. J. M., Deichmann, R., Ashburner, J., & Frackowiak, R. S. J. (2008). Evidence for Segregated and Integrative Connectivity Patterns in the Human Basal Ganglia. Journal of Neuroscience, 28(28), 7143–7152. 10.1523/JNEUROSCI.1486-08.2008

24. Essayed, W. I., Zhang, F., Unadkat, P., Cosgrove, G. R., Golby, A. J., & O’Donnell, L. J. (2017). White matter tractography for neurosurgical planning: A topography-based review of the current state of the art. NeuroImage: Clinical, 15, 659–672. 10.1016/J.NICL.2017.06.011

25. Fischl, B. (2012). FreeSurfer. NeuroImage, 62(2), 774–781. 10.1016/J.NEUROIMAGE.2012.01.021

26. Frigo, M., Deslauriers-Gauthier, S., Parker, D., Ismail, A. A. O., Kim, J. J., Verma, R., & Deriche, R. (2020). Diffusion MRI tractography filtering techniques change the topology of structural connectomes. Journal of Neural Engineering, 17(6), 065002. 10.1088/1741-2552/ABC29B

27. Girard, G., Rafael-Patiño, J., Truffet, R., Aydogan, D. B., Adluru, N., Nair, V. A., Prabhakaran, V., Bendlin, B. B., Alexander, A. L., Bosticardo, S., Gabusi, I., Ocampo-Pineda, M., Battocchio, M., Piskorova, Z., Bontempi, P., Schiavi, S., Daducci, A., Stafiej, A., Ciupek, D., … Thiran, J. P. (2023). Tractography passes the test: Results from the diffusion-simulated connectivity (disco) challenge. NeuroImage, 277, 120231. 10.1016/J.NEUROIMAGE.2023.120231

28. Girard, G., Whittingstall, K., Deriche, R., & Descoteaux, M. (2014). Towards quantitative connectivity analysis: reducing tractography biases. NeuroImage, 98, 266–278. 10.1016/J.NEUROIMAGE.2014.04.074

29. Glasser, M. F., Sotiropoulos, S. N., Wilson, J. A., Coalson, T. S., Fischl, B., Andersson, J. L., Xu, J., Jbabdi, S., Webster, M., Polimeni, J. R., Van Essen, D. C., & Jenkinson, M. (2013). The minimal preprocessing pipelines for the Human Connectome Project. NeuroImage, 80, 105–124. 10.1016/J.NEUROIMAGE.2013.04.127

30. He, Y., Hong, Y., & Wu, Y. (2024). Spherical-deconvolution informed filtering of tractograms changes laterality of structural connectome. NeuroImage, 303, 120904. 10.1016/J.NEUROIMAGE.2024.120904

31. Holm, S. (1979). A Simple Sequentially Rejective Multiple Test Procedure. Scandinavian Journal of Statistics, 6(2), 65–70. http://www.jstor.org/stable/4615733

32. Innocenti, G. M., Dyrby, T. B., Andersen, K. W., Rouiller, E. M., & Caminiti, R. (2017). The Crossed Projection to the Striatum in Two Species of Monkey and in Humans: Behavioral and Evolutionary Significance. Cerebral Cortex, 27(6), 3217–3230. 10.1093/CERCOR/BHW161

33. Innocenti, G. M., Schmidt, K., Milleret, C., Fabri, M., Knyazeva, M. G., Battaglia-Mayer, A., Aboitiz, F., Ptito, M., Caleo, M., Marzi, C. A., Barakovic, M., Lepore, F., & Caminiti, R. (2022). The functional characterization of callosal connections. Progress in Neurobiology, 208, 102186. 10.1016/J.PNEUROBIO.2021.102186

34. Jbabdi, S., & Johansen-Berg, H. (2011). Tractography: Where Do We Go from Here? Brain Connectivity, 1(3), 169–183. 10.1089/BRAIN.2011.0033/ASSET/8A6CDDEA-5A11-48F0-900E-D2C4639F310D/ASSETS/IMAGES/LARGE/10.1089_BRAIN.2011.0033-FIG6.JPG

35. Jeurissen, B., Descoteaux, M., Mori, S., & Leemans, A. (2019). Diffusion MRI fiber tractography of the brain. NMR in Biomedicine, 32(4), e3785. 10.1002/NBM.3785

36. Jeurissen, B., Leemans, A., Tournier, J. D., Jones, D. K., & Sijbers, J. (2013). Investigating the prevalence of complex fiber configurations in white matter tissue with diffusion magnetic resonance imaging. Human Brain Mapping, 34(11), 2747–2766. 10.1002/HBM.22099

37. Jones, D. K., Knösche, T. R., & Turner, R. (2013). White matter integrity, fiber count, and other fallacies: The do’s and don’ts of diffusion MRI. NeuroImage, 73, 239–254. 10.1016/J.NEUROIMAGE.2012.06.081

38. Kellner, E., Dhital, B., Kiselev, V. G., & Reisert, M. (2016). Gibbs-ringing artifact removal based on local subvoxel-shifts. Magnetic Resonance in Medicine, 76(5), 1574–1581. 10.1002/MRM.26054

39. Kita, T., & Kita, H. (2012). The Subthalamic Nucleus Is One of Multiple Innervation Sites for Long-Range Corticofugal Axons: A Single-Axon Tracing Study in the Rat. Journal of Neuroscience, 32(17), 5990–5999. 10.1523/JNEUROSCI.5717-11.2012

40. Koch, P. J., Girard, G., Brügger, J., Cadic-Melchior, A. G., Beanato, E., Park, C. H., Morishita, T., Wessel, M. J., Pizzolato, M., Canales-Rodríguez, E. J., Fischi-Gomez, E., Schiavi, S., Daducci, A., Piredda, G. F., Hilbert, T., Kober, T., Thiran, J. P., & Hummel, F. C. (2022). Evaluating reproducibility and subject-specificity of microstructure-informed connectivity. NeuroImage, 258, 119356. 10.1016/J.NEUROIMAGE.2022.119356

41. Krimmel, S. R., Laumann, T. O., Chauvin, R. J., Hershey, T., Roland, J. L., Shimony, J. S., Willie, J. T., Norris, S. A., Marek, S.N. Van, A., Wang, A., Monk, J., Scheidter, K. M., Whiting, F. I., Ramirez-Perez, N., Metoki, A., Baden, N. J., Kay, B. P., Siegel, J. S., … Dosenbach, N. U. F. (2025). The human brainstem’s red nucleus was upgraded to support goal-directed action. Nature Communications 2025 16:1, 16(1), 3398-. 10.1038/s41467-025-58172-z

42. Lent, R., Tovar-Moll, F., & Szczupak, D. (2026). The ‘secret connections’ of the brain: a connectomic reserve for neuroplasticity? Brain, 139(4), 16–17. 10.1093/BRAIN/AWAG082

43. Leserri, S., & Aydogan, D. B. (2024). HARP: Hierarchical Anatomical Refinement of Pathways in Tractography. International Society of Magnetic Resonance in Medicine (ISMRM) (2024). https://archive.ismrm.org/2024/2160.html

44. Li, C. N., Keay, K. A., Henderson, L. A., & Mychasiuk, R. (2024). Re-examining the Mysterious Role of the Cerebellum in Pain. Journal of Neuroscience, 44(17). 10.1523/JNEUROSCI.1538-23.2024

45. Maffei, C., Girard, G., Schilling, K. G., Aydogan, D. B., Adluru, N., Zhylka, A., Wu, Y., Mancini, M., Hamamci, A., Sarica, A., Teillac, A., Baete, S. H., Karimi, D., Yeh, F. C., Yildiz, M. E., Gholipour, A., Bihan-Poudec, Y., Hiba, B., Quattrone, A., … Yendiki, A. (2022). Insights from the IronTract challenge: Optimal methods for mapping brain pathways from multi-shell diffusion MRI. NeuroImage, 257, 119327. 10.1016/J.NEUROIMAGE.2022.119327

46. Maier-Hein, K. H., Neher, P. F., Houde, J. C., Côté, M. A., Garyfallidis, E., Zhong, J., Chamberland, M., Yeh, F. C., Lin, Y. C., Ji, Q., Reddick, W. E., Glass, J. O., Chen, D. Q., Feng, Y., Gao, C., Wu, Y., Ma, J., Renjie, H., Li, Q., … Descoteaux, M. (2017). The challenge of mapping the human connectome based on diffusion tractography. Nature Communications 2017 8:1, 8(1), 1349-. 10.1038/s41467-017-01285-x

47. Mayberg, H. S. (2009). Targeted electrode-based modulation of neural circuits for depression. The Journal of Clinical Investigation, 119(4), 717. 10.1172/JCI38454

48. Meynert, T., & Sachs, B. (1885). Psychiatry: A Clinical Treatise on Diseases of the Fore-Brain, Based upon a Study of Its Structure, Functions, and Nutrition. Part I: The Anatomy, Physiology, and Chemistry of the Brain: Part I. G. P. Putnam’s Sons.

49. Palesi, F., Tournier, J. D., Calamante, F., Muhlert, N., Castellazzi, G., Chard, D., D’Angelo, E., & Wheeler-Kingshott, C. A. M. (2015). Contralateral cerebello-thalamo-cortical pathways with prominent involvement of associative areas in humans in vivo. Brain Structure and Function, 220(6), 3369–3384. 10.1007/S00429-014-0861-2/FIGURES/6

50. Parent, M., & Parent, A. (2005). Single-axon tracing and three-dimensional reconstruction of centre médian-parafascicular thalamic neurons in primates. Journal of Comparative Neurology, 481(1), 127–144. 10.1002/CNE.20348

51. Petit, L., Mahdy Ali, K., Rheault, F., Boré, A., Cremona, S., Corsini, F., De Benedictis, A., Descoteaux, M., & Sarubbo, S. (2023). The structural connectivity of the human angular gyrus as revealed by microdissection and diffusion tractography. Brain Structure and Function, 228(1), 103–120., *228*, 103–120. 10.1007/s00429-022-02551-5

52. Petit, L., Rheault, F., Descoteaux, M., & Tzourio-Mazoyer, N. (2019). Half of the streamlines built in a whole human brain tractogram is anatomically uninterpretable. *In:* 25th Conference of the Organization for Human Brain Mapping, Roma, 2019. 10.7490/F1000RESEARCH.1118488.1

53. Poulin, P., Theaud, G., Rheault, F., St-Onge, E., Bore, A., Renauld, E., de Beaumont, L., Guay, S., Jodoin, P. M., & Descoteaux, M. (2022). TractoInferno - A large-scale, open-source, multi-site database for machine learning dMRI tractography. Scientific Data 2022 9:1, 9(1), 725-. 10.1038/s41597-022-01833-1

54. Reveley, C., Seth, A. K., Pierpaoli, C., Silva, A. C., Yu, D., Saunders, R. C., Leopold, D. A., & Ye, F. Q. (2015). Superficial white matter fiber systems impede detection of long-range cortical connections in diffusion MR tractography. Proceedings of the National Academy of Sciences of the United States of America, 112(21), E2820–E2828. 10.1073/PNAS.1418198112/SUPPL_FILE/PNAS.201418198SI.PDF

55. Rheault, F., Poulin, P., Caron, A. V., St-Onge, E., Schilling, K. G., Petit, L., Dell’Acqua, F., Leemans, A., & Descoteaux, M. (2025). Current challenges and opportunities for tractography. Handbook of Diffusion MR Tractography: Imaging Methods, Biophysical Models, Algorithms and Applications, 565–580. 10.1016/B978-0-12-818894-1.00037-9

56. Rizzo, M., Luppino, G., & Borra, E. (2026). Qualitative and quantitative analysis of the callosal projections to prefrontal, frontal motor, and parietal areas in the macaque monkey. Brain Structure and Function, 231(1), 3-. 10.1007/S00429-025-03060-X/FIGURES/9

57. Sarwar, T., Ramamohanarao, K., Daducci, A., Schiavi, S., Smith, R. E., & Zalesky, A. (2023). Evaluation of tractogram filtering methods using human-like connectome phantoms. NeuroImage, 281, 120376. 10.1016/J.NEUROIMAGE.2023.120376

58. Schiavi, S., Ocampo-Pineda, M., Barakovic, M., Petit, L., Descoteaux, M., Thiran, J. P., & Daducci, A. (2020). A new method for accurate in vivo mapping of human brain connections using microstructural and anatomical information. Science Advances, 6(31). 10.1126/SCIADV.ABA8245/SUPPL_FILE/ABA8245_SM.PDF

59. Schilling, K. G., Nath, V., Hansen, C., Parvathaneni, P., Blaber, J., Gao, Y., Neher, P., Aydogan, D. B., Shi, Y., Ocampo-Pineda, M., Schiavi, S., Daducci, A., Girard, G., Barakovic, M., Rafael-Patino, J., Romascano, D., Rensonnet, G., Pizzolato, M., Bates, A., … Landman, B. A. (2019). Limits to anatomical accuracy of diffusion tractography using modern approaches. NeuroImage, 185, 1–11. 10.1016/J.NEUROIMAGE.2018.10.029

60. Schilling, K. G., Petit, L., Rheault, F., Remedios, S., Pierpaoli, C., Anderson, A. W., Landman, B. A., & Descoteaux, M. (2020). Brain connections derived from diffusion MRI tractography can be highly anatomically accurate—if we know where white matter pathways start, where they end, and where they do not go. Brain Structure and Function, 225(8), 2387–2402. 10.1007/S00429-020-02129-Z/FIGURES/6

61. Schilling, K., Gao, Y., Janve, V., Stepniewska, I., Landman, B. A., & Anderson, A. W. (2017). Can increased spatial resolution solve the crossing fiber problem for diffusion MRI? NMR in Biomedicine, 30(12), e3787. 10.1002/NBM.3787

62. Schilling, K., Gao, Y., Janve, V., Stepniewska, I., Landman, B. A., & Anderson, A. W. (2018). Confirmation of a gyral bias in diffusion MRI fiber tractography. Human Brain Mapping, 39(3), 1449–1466. 10.1002/HBM.23936

63. Schmahmann, J. D., & Pandya, D. N. (2006). Fiber Pathways of the Brain. Oxford University Press, 1–654. 10.1093/ACPROF:OSO/9780195104233.001.0001

64. Sclocco, R., Beissner, F., Bianciardi, M., Polimeni, J. R., & Napadow, V. (2018). Challenges and opportunities for brainstem neuroimaging with ultrahigh field MRI. NeuroImage, 168, 412–426. 10.1016/J.NEUROIMAGE.2017.02.052

65. Seabold, S., & Perktold, J. (2010). Statsmodels: Econometric and Statistical Modeling with Python. PROC. OF THE 9th PYTHON IN SCIENCE CONF. http://statsmodels.sourceforge.net/

66. Smith, R. E., Tournier, J. D., Calamante, F., & Connelly, A. (2012). Anatomically-constrained tractography: Improved diffusion MRI streamlines tractography through effective use of anatomical information. NeuroImage, 62(3), 1924–1938. 10.1016/J.NEUROIMAGE.2012.06.005

67. Smith, R. E., Tournier, J. D., Calamante, F., & Connelly, A. (2015). SIFT2: Enabling dense quantitative assessment of brain white matter connectivity using streamlines tractography. NeuroImage, 119, 338–351. 10.1016/J.NEUROIMAGE.2015.06.092

68. Smith, S. M., Jenkinson, M., Woolrich, M. W., Beckmann, C. F., Behrens, T. E. J., Johansen-Berg, H., Bannister, P. R., De Luca, M., Drobnjak, I., Flitney, D. E., Niazy, R. K., Saunders, J., Vickers, J., Zhang, Y., De Stefano, N., Brady, J. M., & Matthews, P. M. (2004). Advances in functional and structural MR image analysis and implementation as FSL. NeuroImage, 23(SUPPL. 1), S208–S219. 10.1016/J.NEUROIMAGE.2004.07.051

69. Szczupak, D., Iack, P. M., Rayêe, D., Liu, C., Lent, R., Tovar-Moll, F., & Silva, A. C. (2023). The relevance of heterotopic callosal fibers to interhemispheric connectivity of the mammalian brain. Cerebral Cortex, 33(8), 4752–4760. 10.1093/CERCOR/BHAC377

70. Tang, Y., Sun, W., Toga, A. W., Ringman, J. M., & Shi, Y. (2018). A probabilistic atlas of human brainstem pathways based on connectome imaging data. NeuroImage, 169, 227–239. 10.1016/J.NEUROIMAGE.2017.12.042

71. Thomas, C., Ye, F. Q., Irfanoglu, M. O., Modi, P., Saleem, K. S., Leopold, D. A., & Pierpaoli, C. (2014). Anatomical accuracy of brain connections derived from diffusion MRI tractography is inherently limited. Proceedings of the National Academy of Sciences of the United States of America, 111(46), 16574–16579. 10.1073/PNAS.1405672111/SUPPL_FILE/PNAS.201405672SI.PDF

72. Tournier, J., Calamante, F., & Connelly, A. (2010). Improved probabilistic streamlines tractography by 2nd order integration over fibre orientation distributions. Proceedings of the International Society for Magnetic Resonance in Medicine (2010), p. 1670. https://archive.ismrm.org/2010/1670.html

73. Tournier, J. D., Calamante, F., & Connelly, A. (2012). MRtrix: Diffusion tractography in crossing fiber regions. International Journal of Imaging Systems and Technology, 22(1), 53–66. 10.1002/IMA.22005

74. Tournier, J. D., Smith, R., Raffelt, D., Tabbara, R., Dhollander, T., Pietsch, M., Christiaens, D., Jeurissen, B., Yeh, C. H., & Connelly, A. (2019). MRtrix3: A fast, flexible and open software framework for medical image processing and visualisation. NeuroImage, 202, 116137. 10.1016/J.NEUROIMAGE.2019.116137

75. Tran, G., & Shi, Y. (2015). Fiber Orientation and Compartment Parameter Estimation From Multi-Shell Diffusion Imaging. IEEE Transactions on Medical Imaging, 34(11), 2320–2332. 10.1109/TMI.2015.2430850 10.1109/TMI.2015.2430850

76. Van Essen, D. C., Newsome, W. T., Bixby, J. L., Gordon, H., Hamilton, C., Maunsell, J., & Allman, J. (1982). The pattern of interhemispheric connections and its relationship to extrastriate visual areas in the macaque monkey. Journal of Neuroscience, 2(3), 265–283. 10.1523/JNEUROSCI.02-03-00265.1982

77. Van Essen, D. C., Ugurbil, K., Auerbach, E., Barch, D., Behrens, T. E. J., Bucholz, R., Chang, A., Chen, L., Corbetta, M., Curtiss, S. W., Della Penna, S., Feinberg, D., Glasser, M. F., Harel, N., Heath, A. C., Larson-Prior, L., Marcus, D., Michalareas, G., Moeller, S., … Yacoub, E. (2012). The Human Connectome Project: A data acquisition perspective. NeuroImage, 62(4), 2222–2231. 10.1016/J.NEUROIMAGE.2012.02.018

78. Veraart, J., Fieremans, E., & Novikov, D. S. (2016). Diffusion MRI noise mapping using random matrix theory. Magnetic Resonance in Medicine, 76(5), 1582–1593. 10.1002/MRM.26059

79. Veraart, J., Novikov, D. S., Christiaens, D., Ades-aron, B., Sijbers, J., & Fieremans, E. (2016). Denoising of diffusion MRI using random matrix theory. NeuroImage, 142, 394–406. 10.1016/J.NEUROIMAGE.2016.08.016

80. Wang, R., Chang, Z., Liu, X., Kristanto, D., Gé, É., Gartner, G., Liu, X., Liu, M., Wu, Y., Lui, M., & Zhou, C. (2025). Weak structural connectivity nonlinearly underlying human cognitive abilities. https://arxiv.org/abs/2505.24125v2

81. Warrington, S., Bryant, K. L., Khrapitchev, A. A., Sallet, J., Charquero-Ballester, M., Douaud, G., Jbabdi, S., Mars, R. B., & Sotiropoulos, S. N. (2020). XTRACT - Standardised protocols for automated tractography in the human and macaque brain. NeuroImage, 217, 116923. 10.1016/J.NEUROIMAGE.2020.116923

82. Yamada, K., Sakai, K., Akazawa, K., Yuen, S., & Nishimura, T. (2009). MR Tractography: A Review of Its Clinical Applications. Magnetic Resonance in Medical Sciences, 8(4), 165–174. 10.2463/MRMS.8.165

83. Yeh, C. H., Smith, R. E., Dhollander, T., & Connelly, A. (2017). Mesh-based anatomically-constrained tractography for effective tracking termination and structural connectome construction. Proceedings of the ISMRM (2017), p. 58. https://www.researchgate.net/profile/Chun-Hung-Yeh-3/publication/315836374_Mesh-based_anatomically-constrained_tractography_for_effective_tracking_termination_and_structural_connectome_construction/links/58ee33f6458515c4aa52aebb/Mesh-based-anatomically-constrained-tractography-for-effective-tracking-termination-and-structural-connectome-construction.pdf

84. Yeh, C. H., Smith, R. E., Liang, X., Calamante, F., & Connelly, A. (2016). Correction for diffusion MRI fibre tracking biases: The consequences for structural connectomic metrics. NeuroImage, 142, 150–162. 10.1016/J.NEUROIMAGE.2016.05.047

85. Zalesky, A., Fornito, A., Cocchi, L., Gollo, L. L., van den Heuvel, M. P., & Breakspear, M. (2016). Connectome sensitivity or specificity: which is more important? NeuroImage, 142, 407–420. 10.1016/J.NEUROIMAGE.2016.06.035

